# Oxygen drives a rapid, reversible photoprotective switch in photosynthetic antennae

**DOI:** 10.64898/2026.10.28.762473

**Authors:** Bertus van Heerden, Farooq Kyeyune, Francois Conradie, Sabera Millan, Alastair T. Gardiner, Richard J. Cogdell, Marcin Sarewicz, Artur Osyczka, Michal Gwizdala, Tjaart P. J. Krüger

**Author notes:** These authors contributed equally to this work (Bertus van Heerden, Farooq Kyeyune). Department of Physics, Faculty of Science, Kyambogo University; Kampala, Uganda (Farooq Kyeyune). Department of Chemistry, Ravenshaw University; Cuttack, Odisha, India (Sabera Millan). Institute of Microbiology, Czech Academy of Sciences; Třeboň, Czech Republic (Alastair T. Gardiner). The Institute of Photonic Sciences (ICFO); Barcelona, Spain.

## Abstract

Oxygen is the archetypal agent of photooxidative damage, and it is generally countered by removing its reactive products or by regulated, energy-costly responses. We show that in photosynthetic light-harvesting complexes, oxygen also drives a rapid, reversible photoprotective switch. Using a new bulk fluorescence method that separates reversible dark states from irreversible photobleaching, together with single-molecule spectroscopy, we find that illumination and oxygen generate a reversible, protein-stabilized charge-separated state comprising oxidized chlorophyll and superoxide. Access to this state increasingly outcompetes irreversible photodamage as excitation intensity falls to physiological levels and is strongly accelerated by ascorbic acid in native thylakoid membranes. These findings identify a constitutive oxygen-dependent photoprotective pathway and suggest that redox-state recovery can determine whether oxygen-associated chemistry protects or damages photoactive systems.

---

Illumination can initiate oxygen-dependent photophysical and photochemical pathways, with outcomes ranging from non-destructive excited-state deactivation to the formation of reactive oxygen species (ROS), primarily singlet oxygen and superoxide (O_2_^•−^), that may contribute to irreversible photodamage (*1, 2*). Because fluorescence loss alone cannot distinguish reversible dark-state formation from photobleaching, it remains unclear whether oxygen-dependent pathways can also promote photoprotection. Photosynthetic antennae offer a powerful system in which to test this possibility because their pigment–protein complexes can be studied in isolation and in native membranes, and their established protective pathways provide a physiological benchmark. In these complexes, carotenoids quench chlorophyll (Chl) and bacteriochlorophyll triplet states and singlet oxygen (*3*), various antioxidants remove ROS (*4*), and oxygenic organisms additionally dissipate excess excitation as heat through non-photochemical quenching (NPQ) (*5, 6*). Yet qE, the fastest NPQ component, takes tens of seconds to reach its maximum after irradiance increases (*7–10*), leaving an interval before regulated photoprotection is fully engaged. Moreover, although anoxygenic organisms manage excess excitation and oxidative stress by other means, a rapidly accessible photoprotective process within their antennae has not been established.

The major plant light-harvesting complex (LHCII), the primary site of qE (*11–13*), switches between light-harvesting and quenched states through changes in protein conformation and pigment interactions (*14–16*). Individual complexes also switch between bright and strongly quenched states over milliseconds to minutes (*17, 18*). Because qE-promoting conditions alter this fluorescence blinking favorably, it has been attributed to the conformational dynamics underlying qE (*19, 20*). Yet its molecular origin and physiological significance remain unresolved.

In organic dyes and conjugated polymers, fluorescence blinking involves photoinduced redox reactions with oxygen and the local environment (*21–23*), but a similar mechanism was considered unlikely in photosynthetic complexes because their charge-separated and radical states recombine typically within femtoseconds to microseconds (*24–28*), significantly faster than the millisecond-to-minute OFF states detected by single-molecule spectroscopy (SMS) (*29*). Direct evidence for light-induced radicals in isolated antennae has also been lacking. Thus, whether blinking includes oxygen-dependent photochemistry and can contribute to photoprotection remains unknown.

Assessing photoprotection requires separating dark-state formation, recovery, and cumulative photobleaching at physiological excitation. SMS typically requires intensities far above physiological intensities, while existing fluorescence-fluctuation and transient-state methods do not readily resolve all three processes within one kinetic framework (*30–34*; Text S1). Earlier studies reported oxygen-dependent quenching in aggregated LHCII, but its kinetics in isolated trimers remained unresolved (*35–38*). Here, we introduce Intensity-Programmed Kinetic Isolation (IPKI) and combine it with SMS to connect ensemble kinetics with single-complex blinking. Oxygen manipulation, superoxide-sensitive probes, and separate EPR measurements identify superoxide-linked oxidation and light-induced radicals. We identify oxygen-dependent reversible cycling in isolated LHCII, bacterial LH2 and native thylakoids, supporting a photoprotective role.

## IPKI separates reversible dark-state cycling from photobleaching

IPKI extends pulse-induced relaxation measurements by independently controlling the excitation used to prepare photophysical states and the illumination used to read their recovery (Fig. 1A). During each cycle, a high-intensity preparation interval drives the population into long-lived dark states, after which partial recovery is allowed during a dark period. The fluorescence intensity is monitored during the light and dark phases using low-intensity pulses that minimally perturb the system. Repeating the cycle produces a bulk fluorescence trace in which reversible recovery and cumulative photobleaching can be quantified within a common kinetic framework from sub-milliseconds to tens of minutes (Fig. 1B). Analyzing individual cycles also allows the model to capture kinetic heterogeneity.

**Fig 1.**
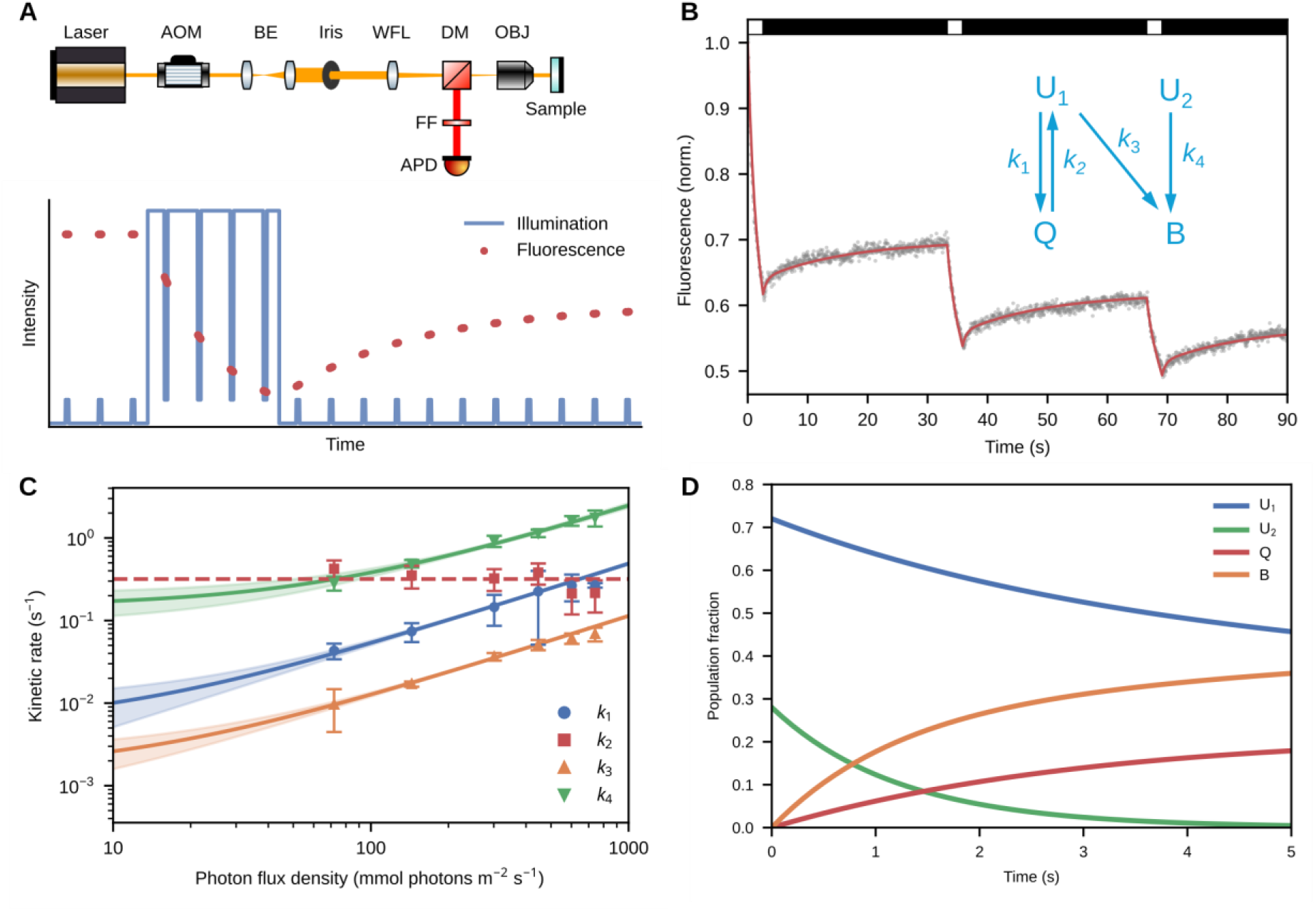
IPKI resolves reversible dark-state dynamics and photobleaching. (**A**) Top: Schematic of IPKI setup. AOM: acousto-optic modulator, BE: beam expander, WFL: wide-field lens, DM: dichroic mirror, OBJ: objective lens, FF: fluorescence filter, APD: avalanche photodiode. See Materials and Methods for more details. Bottom: Illustration of IPKI principle: the baseline fluorescence intensity is first recorded, high illumination then populates dark states, after which a dark period enables fluorescence recovery. Short, low-intensity pulses are used to probe the fluorescence. (**B**) Example of experimental IPKI trace measured on isolated LHCII. Black and white bars on top indicate low- and high-illumination intensities, respectively. Grey: data; red: fit. Inset: effective kinetic model used to fit the data, showing the pooled reversible quenched-state (Q) population and effective rates *k*_1_ and *k*_2_. Actual fits were done using the extended model in Figure S1D to capture the biphasic recovery of Q. See Materials and Methods for details. (**C**) Excitation-intensity dependence of effective kinetic rates determined from global IPKI analysis, shown on a double logarithmic scale, with linear fits (lines) and corresponding 68% confidence intervals (shaded). The recovery rate *k*_2_ was independent of excitation intensity and is shown as a constant value (dashed line). The corresponding analysis allowing light-dependent recovery is shown in Figure S2A; fitting details are described in the Materials and Methods. Errors denote Jackknife 95% confidence intervals. Plots on a linear scale are shown in Figure S3. (**D**) Simulated population kinetics for LHCII illuminated continuously with 300 mmol photons m^−2^ s^−1^. Population fractions for the extended kinetic model and for the accumulated Q population are shown in Figure S4.

The minimal phenomenological model comprises two fluorescent populations, U_1_ and U_2_, a reversible dark state Q, and an irreversibly bleached state B (Fig. 1B, inset). The effective rates *k*_1_ and *k*_2_ describe population of and recovery from Q, respectively, and *k*_3_ and *k*_4_ describe bleaching from U_1_ and U_2_, respectively. Removing U_2_ produced strong, systematic deviations from the experimental transients (Fig. S1B), while for LHCII, biexponential recovery noticeably improved the fits, which was represented by two effective recovery components, Q_1_ and Q_2_ (Fig. S1D).

Across the measured excitation range, *k*_1_, *k*_3_, and *k*_4_ increase approximately linearly with photon flux, whereas the recovery components (*k*_2a_ and *k*_2b_, and hence *k*_2_) do not (Fig. 1C, Fig. S2A, and Table S1). For LHCII, the regression of *k*_2_ against excitation intensity gave *p* = 0.09, and a model allowing additional light-dependent recovery fitted less well, with a Bayesian information criterion of −104 compared with −118 for the intensity-independent model. These results provide no statistically significant evidence for a linear intensity dependence of recovery (Text S2).

The fitted *k*_1_ exceeded *k*_3_ throughout the measured range, favoring entry into the reversible dark state over direct bleaching from U_1_. U_2_ denotes a small, highly photolabile population that dominated the initial fluorescence decrease but was largely depleted within the first seconds of illumination (Fig. 1D). As excitation intensity decreased, the light-driven transition and bleaching rates slowed, while the recovery rate remained intensity-independent. This is also reflected by the linear dependence of the reversible dark-state occupancy at low excitation intensities (Fig. S5A). The resulting kinetics support a rapidly accessible reversible pathway that can contribute to photoprotection at low excitation intensities, particularly before regulated NPQ is fully established.

### Oxygen controls reversible switching in LHCII and LH2

An oxygen-scavenging system (glucose, glucose oxidase, and catalase; GOC) substantially decreased *k*_1_, *k*_3_, and *k*_4_ in LHCII (Fig. 2A and Table S1), demonstrating that both reversible dark-state formation and irreversible bleaching depend on molecular oxygen. In contrast, recovery (*k*_2_) increased through its fast component, *k*_2a_, whereas the slow component, *k*_2b_, decreased slightly (Table S1), indicating that the recovery pathways within the heterogeneous dark-state manifold differ in oxygen sensitivity.

**Fig 2.**
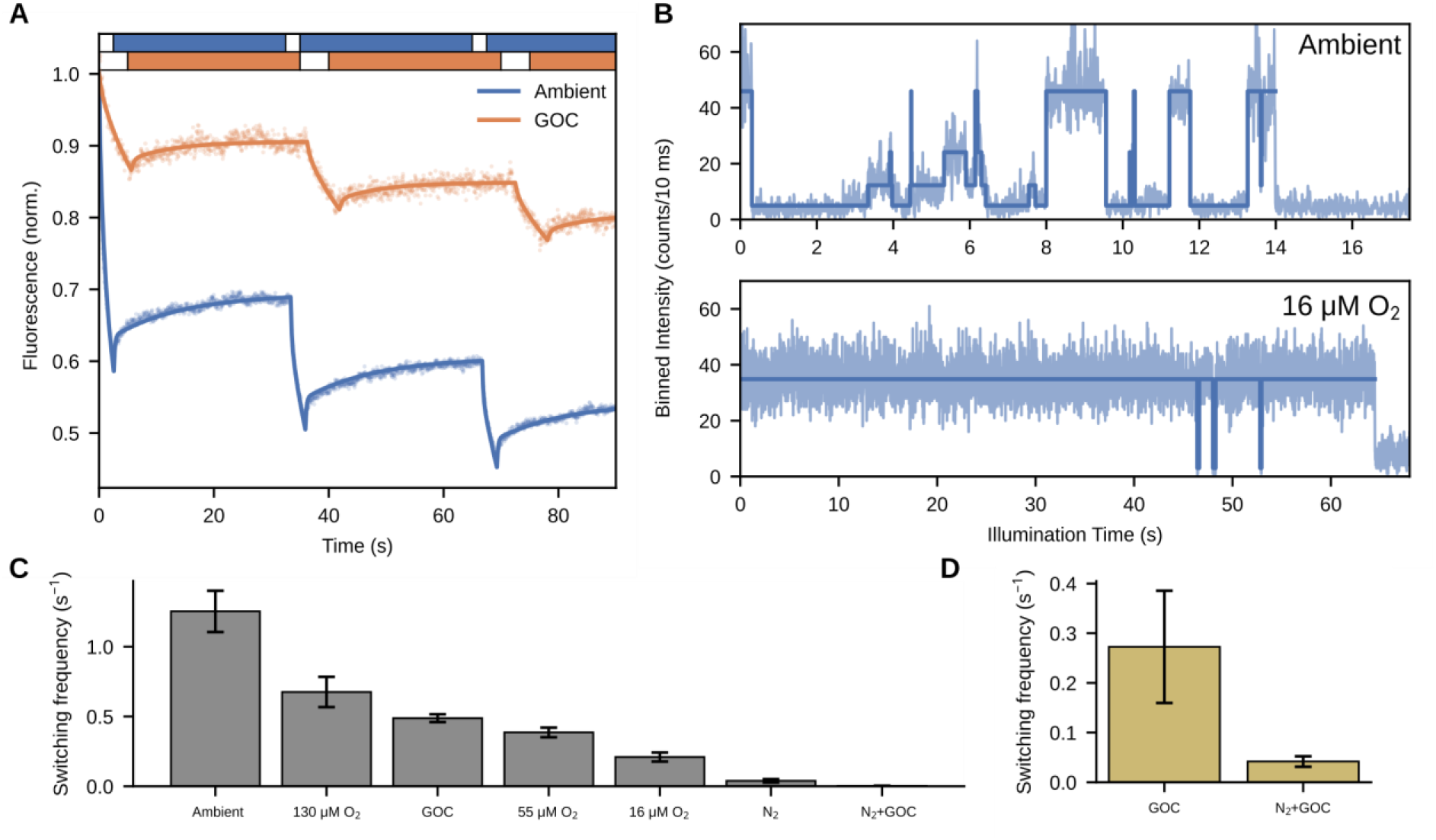
Oxygen dependence of reversible quenching in LHCII and LH2. (**A**) IPKI traces for LHCII with and without GOC at 301 mmol photons m^−2^ s^−1^, using a two-times longer illumination time for the GOC condition, which denotes enzymatic reduction of oxygen through the addition of a glucose–glucose oxidase–catalase mixture to the sample. Bars on top indicate alternating periods of high and low illumination. (**B**) Example intensity traces of single LHCII trimers in ambient conditions (270 μM O_2_, top) and at 16 μM O_2_ (bottom), showing significantly reduced blinking and bleaching at a lower oxygen concentration. Binned data (light blue) were fitted and grouped into binary (ON and OFF) states – see Materials and Methods. (**C**) Average switching frequency between ON and OFF states of single LHCII complexes under different oxygen conditions. Dissolved oxygen concentrations followed from continuous flushing of the sample cell by O_2_/N_2_ gas mixtures and were calculated from the corresponding gas-phase oxygen fractions assuming air–water equilibrium under the experimental conditions. N_2_ denotes continuous flushing of the sample cell with pure N_2_ gas. The gas-flushing setup is depicted in Figure S6. GOC reduced the effective dissolved oxygen concentration to ∼90 μM, equivalent to equilibration with a gas phase containing ∼7% O_2_. (**D**) Average switching frequency between ON and OFF states of single LH2 complexes at different oxygen conditions. Error bars denote standard errors.

SMS provided an independent test of the oxygen dependence of fluorescence switching. As oxygen concentration decreased, switching between emissive and quenched levels became progressively less frequent (Fig. 2, B and C). Nitrogen purging strongly suppressed switching, and the combination of nitrogen purging and GOC nearly abolished detectable events. The close correspondence between oxygen-dependent changes in bulk IPKI kinetics and single-complex switching supports a link between the reversible dark-state process resolved by IPKI and fluorescence blinking observed by SMS. Blinking was also observed in freely diffusing LHCII (Fig. S5D), showing that surface immobilization does not cause it.

A similar oxygen dependence was observed in single LH2 complexes from the purple bacterium *Rhodoblastus acidophilus*, an anoxygenic antenna system that lacks plant-type qE (Fig. 2D). Earlier SMS studies of LH2 reported an excitation-dependent quenching rate and a recovery rate largely independent of excitation intensity (*40*), as we find for LHCII (Fig. 1). The results therefore establish oxygen-dependent reversible switching in representative oxygenic and anoxygenic antenna complexes, suggesting a broader photoprotective principle of photosynthetic antennae. Because anoxygenic organisms can encounter oxygen despite living in low-oxygen environments, a constitutive pathway of this kind could protect them during transient oxidative stress even without plant-type qE.

### Superoxide and a radical accompany the dark state

To investigate whether oxygen-dependent dark-state formation involves Type I photooxidative chemistry, we tested for 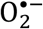-associated oxidation using dihydroethidium (DHE). Strong illumination of LHCII produced a broad emission band near 620 nm that was strongly suppressed by superoxide dismutase (SOD; Fig. 3A). This response is consistent with 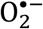-dependent formation of DHE oxidation products. Because DHE can undergo multiple oxidation reactions, its fluorescence does not uniquely identify 2-hydroxyethidium or quantify 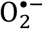 selectively (*41*).

**Fig 3.**
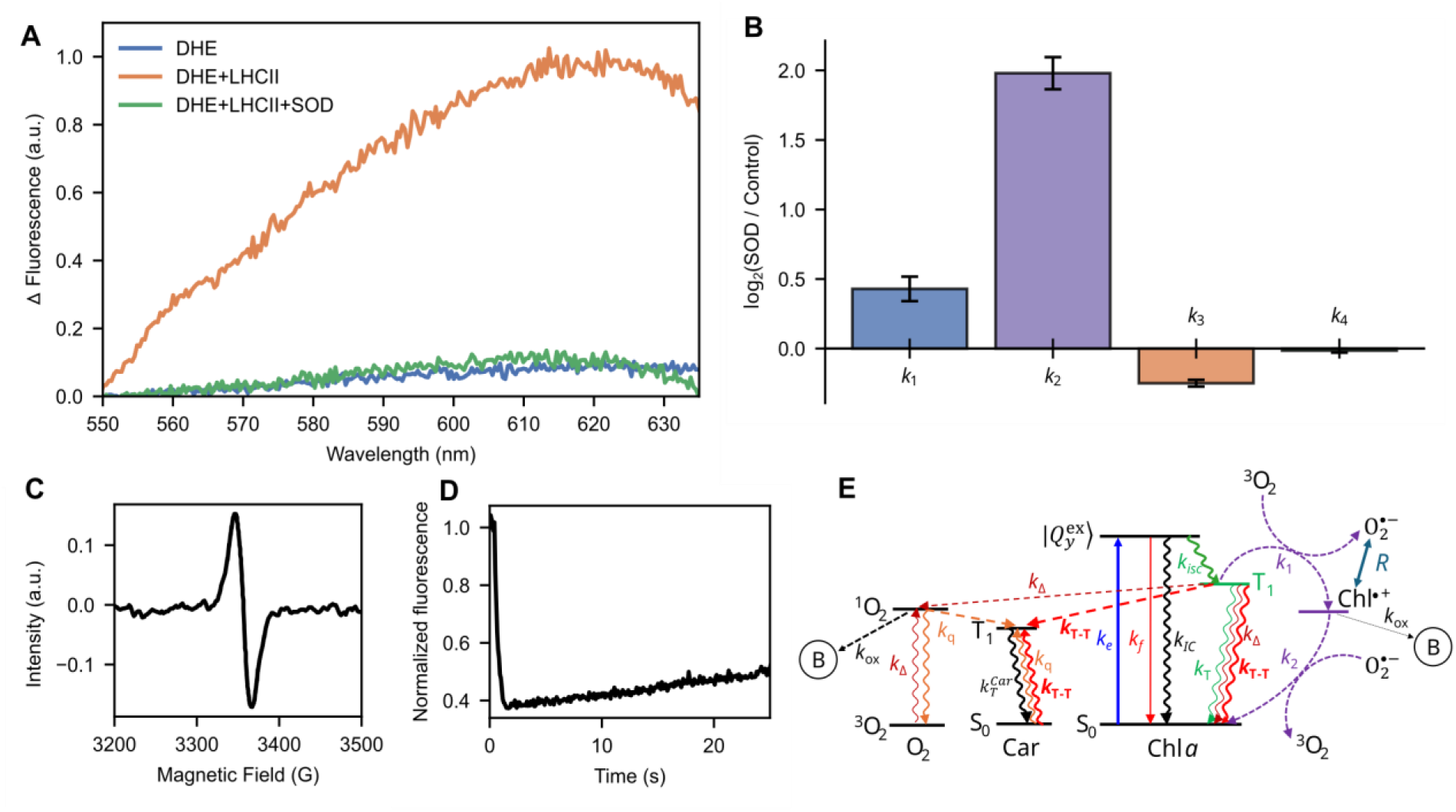
Superoxide, a light-induced radical and the requirement for the protein environment. (**A**) Difference fluorescence spectra after 4 min of strong illumination of LHCII mixed with DHE (orange) and compared with a DHE solution (blue) and LHCII mixed with both DHE and SOD (green) illuminated under the same conditions. The weak emission from the DHE solution is consistent with low-level formation of DHE oxidation products. The spectra at the start and end of the illumination period are shown in Figure S7. (**B**) Log_2_ fold change in IPKI kinetic parameters upon addition of SOD to LHCII. The control depicts the measurement without SOD. The corresponding IPKI traces, the raw spectra, and the values for *k*_2a_ and *k*_2b_ are shown in Figure S7. Errors denote standard errors. (**C**) Electron paramagnetic resonance (EPR) spectrum of LHCII at 15 K under strong illumination, indicating the accumulation of a light-induced organic radical. (**D**) IPKI trace measured for Chl *a* in a 60% ethanol / 40% glycerol (v/v) solution. (**E**) Proposed photophysical and photochemical pathways for oxygen-dependent quenching and competing processes.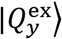: lowest *Q*_*y*_ exciton state of LHCII, *R*: conformational coordinate gating the interaction between the two radicals, Car: carotenoid, ^3^O_2_: ground-state molecular oxygen, *k*_1_ (*k*_2_): phenomenological kinetic rate associated with dark-state (de)population, regulated by radical-pair formation (recombination) and protein conformational gating, rates of competing processes: *k*_*e*_: optical excitation, *k*_*f*_: fluorescence, *k*_*IC*_: internal conversion, *k*_*isc*_: intersystem crossing, *k*_T_: intrinsic ^3^Chl decay, *k*_Δ_: singlet oxygen generation (Type II photosensitization), *k*_T-T_: triplet-triplet energy transfer, *k*_*q*_: singlet oxygen quenching by Car, 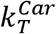: intrinsic ^3^Car decay, *k*_ox_: oxidative/photochemical damage.

We next tested how SOD affected the kinetics resolved by IPKI. SOD increased *k*_1_ by ≈35% and the average recovery rate *k*_2_ about fourfold, decreased *k*_3_ by ≈16%, and left *k*_4_ essentially unchanged (Fig. 3B). The pronounced acceleration of recovery implicates an SOD-sensitive, 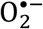-linked redox process in the reversible dark state kinetics. The increase in *k*_1_ likely reflects oxygen regenerated by O_2_^•−^ dismutation, although other changes in the redox environment cannot be excluded. The much larger increase in *k*_2_ shows that removing accessible O_2_^•−^ strongly accelerates recovery, and the small change in *k*_3_ and none in *k*_4_ suggest that O_2_^•−^ is not required for the dominant irreversible pathway. These results support a direct role for O^2•−^ in the formation and recovery of the quenching state.

EPR of LHCII at 15 K independently showed a narrow light-induced signal at 3356 G (Fig. 3C), indicating accumulation of an organic radical whose resonance is compatible with a Chl radical cation.

### The protein stabilizes the charge-separated state

How can an oxygen-dependent charge-separated state last for seconds when photoinduced charge-separated and radical states in photosynthetic complexes recombine much faster (*27, 28*)? Even in reaction centers, where dedicated cofactors stabilize charge separation, recombination is orders of magnitude faster than the dark states seen here (*24–26*). We therefore asked whether the protein environment is required to stabilize the dark states. IPKI of freely diffusing Chl *a* in ethanol showed no fluorescence recovery beyond ∼10 ms (Fig. 3D); the slow rise at later times reflects diffusion of molecules into the observation volume (Fig. S7G). Thus, isolated Chl *a* has no detectable long-lived reversible dark state, indicating that the protein environment is essential for stabilizing the quenched state in LHCII. The intensity-independent recovery (Fig. 1) and the power-law OFF-state statistics reported previously (*18*) are consistent with slow conformational dynamics contributing to recovery.

The oxygen dependence, DHE/SOD response, light-induced radical signal, and requirement for a protein environment are most consistent with an oxygen-dependent charge-separated state involving oxidized Chl and O_2_^•−^. We therefore propose the mechanistic model in Figure 3E in which a Chl triplet transfers an electron to molecular oxygen, generating a Chl radical cation (Chl^•+^) and O_2_ ^•−^. The protein transiently stabilizes this pair, with conformational fluctuations modulating the electronic coupling between the charges. Recovery occurs when thermally activated structural fluctuations restore favorable electronic coupling and permit charge recombination. In this model, *k*_1_ represents formation of the charge-separated state, *k*_2_ its conformationally gated recovery, and *k*_3_ and *k*_4_ competing irreversible photochemical pathways. The hole may be delocalized over several coupled Chls and possibly neighboring carotenoids, which could lower the free energy of the pair and add to its stability.

### Reversible quenching in native membranes

To test whether the mechanism operates in a native membrane under physiologically relevant conditions, we applied IPKI to isolated thylakoid membranes over a wide range of intensities extending into the physiological range, with and without ascorbic acid (AA). The recovery rate *k*_2_ was again independent of excitation intensity (*p* = 0.72 without AA and 0.29 with AA; Figs. 4A and S2B), whereas the photoinduced rates retained their intensity dependence. A single recovery component sufficed, unlike the biexponential recovery in isolated LHCII. Across most intensities, the rates exceeded those of LHCII (Fig. 4A and Table S3). As in LHCII, *k*_1_ exceeded *k*_3_, and *k*_2_ exceeded the photoinduced rates below ≈100 mmol photons m^−2^ s^−1^, indicating that recovery from the dark state is increasingly favored over bleaching as excitation intensity decreases.

**Fig 4.**
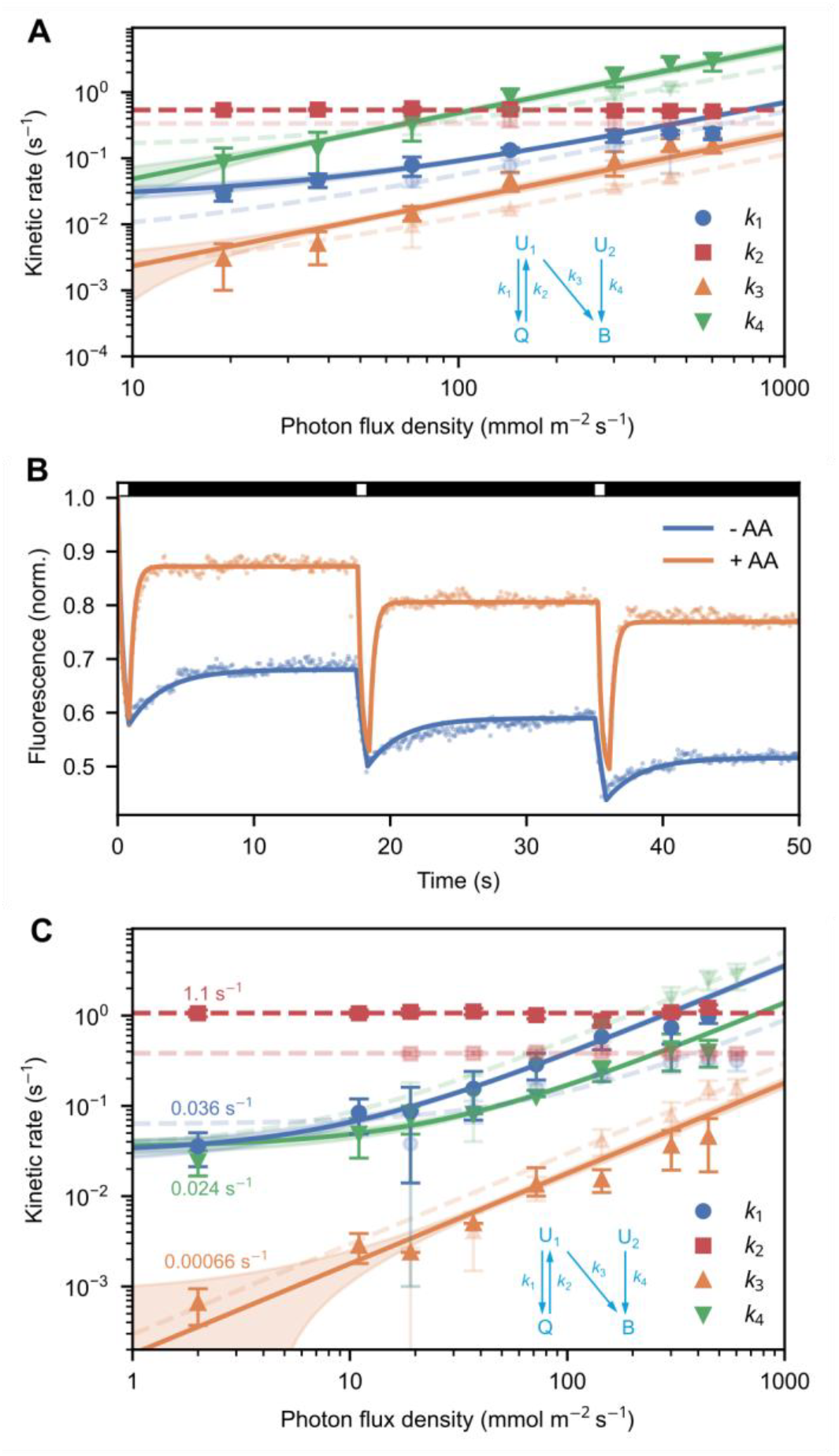
Reversible quenching in thylakoid membranes. (**A**) Power dependence of kinetic rates determined from global IPKI analysis for intact thylakoids in the absence of ascorbic acid (AA), based on the kinetic model in the inset. The recovery rate *k*_2_ was independent of excitation intensity and is shown as a constant value (dashed line). The corresponding analysis allowing light-dependent recovery is shown in Figure S2B. Linear fits (lines) are shown with corresponding 68% uncertainty intervals (shaded). Data for LHCII are shown in a corresponding transparent color. Errors denote Jackknife 95% confidence intervals. Linear plots are shown in Figure S3. (**B**) Representative IPKI traces of thylakoids showing the effect of AA (orange) compared to the absence of AA (blue): a reduction in bleaching and enhancement in reversible quenching. Fits (lines) are shown over data points. Black and white bars on top indicate low and high illumination intensities, respectively. (**C**) Power dependence of kinetic rates for thylakoids with AA present, based on the kinetic model in the inset. *k*_2_ was treated as for the other samples. Linear fits (lines) are shown with corresponding 68% uncertainty intervals (shaded). Data for thylakoids without AA are shown in a corresponding transparent color. Values for kinetic rates measured at 2,000 μmol photons m^−2^ s^−1^ are indicated. Errors denote Jackknife 95% confidence intervals. Linear plots are shown in Figure S3.

AA strongly accelerated recovery from the dark state and suppressed the competing irreversible pathways (Fig. 4, B and C). The reduction in bleaching agrees with the established antioxidant function of ascorbate in chloroplasts, and the increase in *k*_2_ indicates that reduction of the photooxidized state strongly favors recovery of the fluorescent state, lowering the probability of bleaching that involves Chl^•+^ or O_2_ ^•−^. AA also increased *k*_1_, robustly to changes in the fitted *k*_2_ onstraint (Fig. S5C), possibly through changes in the local redox environment or triplet-state kinetics. The strong acceleration of recovery observed independently through *k*_2_ supports enhanced recovery as a major effect of AA. A small deviation at the highest intensities likely reflects singlet–triplet annihilation (*42*). Because this effect is confined to the highest excitation regime and does not alter our conclusions concerning the excitation dependence of dark-state formation and bleaching or the excitation independence of recovery, it was not incorporated into the kinetic model.

Strikingly, *k*_1_ exceeded *k*_3_ at all intensities, by about an order of magnitude at the highest intensities, with the ratio increasing progressively as excitation decreased (Fig. S5C). Because *k*_1_ and *k*_3_ are competing transitions from U_1_, this increasing ratio indicates that entry into the reversible dark state becomes increasingly favored over direct bleaching from U_1_ at lower excitation. Meanwhile, *k*_2_ remains approximately intensity-independent, allowing the quenched population to recover rapidly as the light-driven rates decrease. The alternative bleaching rate *k*_4_ approached *k*_1_ at the lowest intensities but acts on the small, photolabile U_2_ population, which is depleted early during illumination (Fig. S4, C and D). Together, these kinetic relationships indicate a balance strongly shifted toward rapidly reversible Q state population as excitation decreases, providing a quantitative basis for the photoprotective role of the oxygen-dependent pathway at physiological intensities. Ascorbate concentrations in chloroplasts are substantial, rising particularly during high-light acclimation (*43*), suggesting that the redox-sensitive pathway identified here could be even more enhanced *in vivo*.

### Blinking, qE, and photoprotection

Our results fundamentally revise the interpretation of fluorescence blinking in photosynthetic antenna complexes. Previous studies considered blinking to be a direct reporter of the conformational dynamics underlying activated, regulated photoprotection (*19, 39, 44–46*), and this view has been extended to antennae of other oxygenic organisms (*47–49*). We find instead that the long-lived dark states arise from an oxygen-dependent photochemical cycle in which protein conformational fluctuations modulate the formation, stabilization, and recombination of a reversible charge-separated state. This cycle is chemically and kinetically distinct from qE, a regulated process that requires lumen acidification, PsbS, and xanthophyll-cycle activation. It needs no upstream signal, feedback loop, or coordinated activation step, it operates whenever excitation and oxygen are present, and occurs in LH2, which lacks plant-type qE. Blinking is therefore not intrinsically a manifestation of qE but reflects a more general property of photosynthetic antenna photophysics.

Faster dark-state recovery in thylakoids shows that the membrane environment tunes the dark-state kinetics, possibly through pigment–protein interactions, lipid organization, local redox conditions, and/or neighboring complexes. Because the same kinetics occur in isolated LHCII, the mechanism originates within the antenna and does not require reaction-center photochemistry.

Earlier studies found oxygen-dependent, light-induced quenching in aggregated LHCII that recovers over tens of minutes to hours, but not in trimers (*35, 36, 38*). Our results show that the underlying oxygen-dependent dark-state cycle can also occur in individual complexes, although with substantially faster recovery (10–20 s). Thus, aggregation can enhance and prolong the process but is not required. Whether the aggregation-induced effects arise from enhanced excitonic connectivity, changes in pigment–protein organization, or both remains to be determined.

The Chl^•+^–O_2_^•−^ pairing remains a mechanistic model because, although the EPR data are consistent with a Chl-type radical cation, they do not show its pairing with O_2_^•−^ or locate the positive charge, and DHE reports only the accessible O_2_^•−^ pool. The selective SOD sensitivity of *k*_2a_ suggests that the two recovery components sample environments with different access to aqueous O_2_^•−^. Likewise, U_2_ is a kinetic, not structural, assignment, and may represent transient heterogeneity or additional photochemical pathways not included in the minimal model.

The framework also resolves an apparent tension between ultrafast and single-molecule studies: the OFF-state lifetime need not correspond to that of an isolated radical pair but can report a protein-stabilized configuration in which structural fluctuations gate recombination. Long-lived Chl^•+^ is not unprecedented in photosynthetic proteins, as shown by the slow recombination of Chl_Z_ in the Photosystem II reaction center under conditions that restrict conventional electron-transfer pathways (*50–52*).

The mechanism resolved here does more than quench fluorescence: it provides a reversible route to temporarily divert excitation into a redox state and return it to the light-harvesting manifold before irreversible damage occurs, thus playing a photoprotective function. Because the pathway is constitutive, it is well suited to the interval after a sudden rise in irradiance, before regulated NPQ responses become fully established.

### A broader principle

Oxygen-dependent charge transfer can lead either to reversible photoprotection or to irreversible photodamage. The outcome depends on whether both charge species can be safely neutralized or recombine before damaging reactions accumulate. Photosynthetic antennae are embedded in a protective redox environment: carotenoids quench reactive excited states and participate in radical chemistry, while ascorbate and other antioxidants provide additional means of managing oxidized species and ROS. Together with the protein environment stabilizing the charge-separated state, these defenses may favor recovery of the photoactive state over irreversible oxidation.

Many other biological pigment systems likewise operate within cellular antioxidant and redox-regulatory networks. By contrast, many artificial photoactive materials lack similar integrated mechanisms for charge neutralization, damage prevention, and molecular repair. In poly(3-hexylthiophene), for example, light-driven oxygen reduction and hydrogen peroxide production accompany irreversible polymer photooxidation, even when adding sacrificial electron donors (*53*). This illustrates how oxygen-dependent redox chemistry can consume a photoactive material instead of sustaining its operation.

This comparison suggests a broader principle: the consequences of oxygen-dependent redox chemistry depend not simply on charge separation or ROS formation, but on the molecular environment’s capacity to manage the resulting oxidized and reduced species and restore the photoactive state before irreversible reactions dominate. Testing how broadly this principle applies will require examining biological pigments and synthetic materials with different redox partners, antioxidant capacity, and mechanisms of molecular repair. Comparing antennae with different triplet yields and pigment organizations could help establish the role of the proposed triplet precursor. Perturbing the carotenoid content or antioxidant availability could test whether weakening charge neutralization shifts the balance from reversible cycling toward bleaching. Other biological pigments, such as melanin, and synthetic photoactive materials provide further systems to investigate the kinetic principles.

IPKI offers a way to resolve reversible and irreversible processes in fluorophores, quantum emitters, and light-harvesting materials. It may also help characterize dark-state recovery and photobleaching in photosensitizers, providing a kinetic framework for investigating how their photostability or phototoxicity can be optimized. More broadly, these findings suggest that controlling the fate of photoinduced redox states—not simply preventing oxygen-dependent chemistry—could provide a design principle for natural and artificial light-driven systems.

## Acknowledgments

We thank Robert Ekiert for assistance with the EPR measurements.

## Funding

National Research Foundation, South Africa, grant 102431 (FK)

National Research Foundation, South Africa, grant 137973 (SM, TPJK)

National Research Foundation, South Africa, grants 109302, 112085, 150731 and CPRR240403211945 (TPJK)

University of Pretoria postdoctoral fellowships (BvH, FK, MG)

South African Quantum Technology Initiative (BvH, FC)

Laser Rental Pool Program of the Photonics Centre, Council for Scientific and Industrial Research, South Africa (TPJK)

Biotechnology and Biological Sciences Research Council grant BB/N016734/1 (RJC, ATG)

Photosynthetic Antenna Research Center, an Energy Frontier Research Center funded by the U.S. Department of Energy, Office of Science, Office of Basic Energy Sciences, award DE-SC0001035 (RJC, ATG)

Strategic Program Excellence Initiative at Jagiellonian University, BioS PRA (AO, MS)

## Author contributions

Conceptualization: TPJK, MG, FK

Methodology: BvH, FC

Investigation: BvH, FC, FK, MG, MS, AO

Resources: MG, BvH, SM, FC, ATG, RJC

Funding acquisition: TPJK

Project administration: TPJK

Formal analysis: BvH, TPJK

Supervision: TPJK

Writing – original draft: TPJK

Writing – review & editing: BvH, FK, FC, SM, ATG, RJC, MS, AO, MG, TPJK

## Supplementary Materials

Materials and Methods

Supplementary Text

Figs. S1 to S7

Tables S1 to S4

## References

1. X.-D. Wang, O. S. Wolfbeis, Optical methods for sensing and imaging oxygen: Materials, spectroscopies and applications. Chem. Soc. Rev. 43, 3666–3761 (2014).

2. M. S. Baptista, J. Cadet, A. Greer, A. H. Thomas, Photosensitization reactions of biomolecules: Definition, targets and mechanisms. Photochem. Photobiol. 97, 1456–1483 (2021).

3. A. J. Young, H. A. Frank, Energy transfer reactions involving carotenoids: quenching of chlorophyll fluorescence. J. Photochem. Photobiol. B Biol. 36, 3–15 (1996).

4. A. Krieger-Liszkay, Singlet oxygen production in photosynthesis. J. Exp. Bot. 56, 337–346 (2005).

5. B. Demmig-Adams, Carotenoids and photoprotection in plants: a role for the xanthophyll zeaxanthin. Biochim. Biophys. Acta Bioenerg. 1020, 1–24 (1990).

6. H. van Amerongen, R. Croce, Non-photochemical quenching in plants: mechanisms and mysteries. Plant Cell koaf240 (2025).

7. L. A. I. Ramakers, J. Harbinson, E. Wientjes, H. van Amerongen, Unravelling the different components of nonphotochemical quenching using a novel analytical pipeline. New Phytol. 245, 625–636 (2025).

8. L. A. I. Ramakers, J. Harbinson, H. van Amerongen, A novel multivariate analysis: overturning long-held beliefs about non-photochemical quenching. Physiol. Plant. 177, e70420 (2025).

9. L. Lam et al., Dissecting the contributions to non-photochemical quenching in a land plant under fluctuating light. Nat. Commun. 17, 3664 (2026).

10. A. Derks, K. Schaven, D. Bruce, Diverse mechanisms for photoprotection in photosynthesis. Dynamic regulation of photosystem II excitation in response to rapid environmental change. Biochim. Biophys. Acta Bioenerg. 1847, 468–485 (2015).

11. Z. Guardini et al., Mapping light-harvesting and photoprotection responses in the Photosystem II antenna system of higher plants. Plant Physiol. 199, kiaf588 (2025).

12. L. Nicol, W. J. Nawrocki, R. Croce, Disentangling the sites of non-photochemical quenching in vascular plants. Nat. Plants 5, 1177–1183 (2019).

13. L. Nicol et al., The quantitative contribution of different Photosystem II compartments to non-photochemical quenching in Arabidopsis. Nat. Commun. 17, 8051 (2026).

14. A. A. Pascal et al., Molecular basis of photoprotection and control of photosynthetic light-harvesting. Nature 436, 134–137 (2005).

15. V. Daskalakis, S. Papadatos, U. Kleinekathöfer, Fine tuning of the photosystem II major antenna mobility within the thylakoid membrane of higher plants. Biochim. Biophys. Acta Biomembr. 1861, 183059 (2019).

16. M. Ruan et al., Cryo-EM structures of LHCII in photo-active and photo-protecting states reveal allosteric regulation of light harvesting and excess energy dissipation. Nat. Plants 9, 1547–1557 (2023).

17. T. P. J. Krüger, C. Ilioaia, R. van Grondelle, Fluorescence intermittency from the main plant light-harvesting complex: resolving shifts between intensity levels. J. Phys. Chem. B 115, 5071–5082 (2011).

18. T. P. J. Krüger, C. Ilioaia, L. Valkunas, R. van Grondelle, Fluorescence intermittency from the main plant light-harvesting complex: sensitivity to the local environment. J. Phys. Chem. B 115, 5083–5095 (2011).

19. T. P. J. Krüger et al., Controlled disorder in plant light-harvesting complex II explains its photoprotective role. Biophys. J. 102, 2669–2676 (2012).

20. L. Valkunas, J. Chmeliov, T. P. J. Krüger, C. Ilioaia, R. van Grondelle, How photosynthetic proteins switch. J. Phys. Chem. Lett. 3, 2779–2784 (2012).

21. R. Zondervan, F. Kulzer, S. B. Orlinskii, M. Orrit, Photoblinking of rhodamine 6G in poly(vinyl alcohol): radical dark state formed through the triplet. J. Phys. Chem. A 107, 6770–6776 (2003).

22. J. Yu, D. Hu, P. F. Barbara, Unmasking electronic energy transfer of conjugated polymers by suppression of O2 quenching. Science 289, 1327–1330 (2000).

23. T. Christ, F. Kulzer, P. Bordat, T. Basché, Watching the photo-oxidation of a single aromatic hydrocarbon molecule. Angew. Chem. Int. Ed. 40, 4192–4195 (2001).

24. A. R. Holzwarth et al., Kinetics and mechanism of electron transfer in intact photosystem II and in the isolated reaction center: pheophytin is the primary electron acceptor. Proc. Natl. Acad. Sci. U.S.A. 103, 6895–6900 (2006).

25. Y. Yoneda et al., The initial charge separation step in oxygenic photosynthesis. Nat. Commun. 13, 2275 (2022).

26. M. Y. Dekmak, S. M. Mäusle, J. Brandhorst, P. S. Simon, H. Dau, Tracking the first electron transfer step at the donor side of oxygen-evolving photosystem II by time-resolved infrared spectroscopy. Photosynth. Res. 162, 353–369 (2024).

27. N. E. Holt et al., Carotenoid cation formation and the regulation of photosynthetic light harvesting. Science 307, 433–436 (2005).

28. T. K. Ahn et al., Architecture of a charge-transfer state regulating light harvesting in a plant antenna protein. Science 320, 794–797 (2008).

29. J. M. Gruber, P. Malý, T. P. J. Krüger, R. van Grondelle, From isolated light-harvesting complexes to the thylakoid membrane: a single-molecule perspective. Nanophotonics 7, 81–92 (2018).

30. J. Widengren, R. Rigler, Mechanisms of photobleaching investigated by fluorescence correlation spectroscopy. Bioimaging 4, 149–157 (1996).

31. A. Ghosh, J. Enderlein, Advanced fluorescence correlation spectroscopy for studying biomolecular conformation. Curr. Opin. Struct. Biol. 70, 123–131 (2021).

32. T. Schröder et al., Shrinking gate fluorescence correlation spectroscopy yields equilibrium constants and separates photophysics from structural dynamics. Proc. Natl. Acad. Sci. U.S.A. 120, e2211896120 (2023).

33. J. Widengren, Fluorescence-based transient state monitoring for biomolecular spectroscopy and imaging. J. R. Soc. Interface 7, 1135–1144 (2010).

34. J. Tornmalm, J. Widengren, Label-free monitoring of ambient oxygenation and redox conditions using the photodynamics of flavin compounds and transient state (TRAST) spectroscopy. Methods 140–141, 178–187 (2018).

35. R. C. Jennings, F. M. Garlaschi, G. Zucchelli, Light-induced fluorescence quenching in the light-harvesting chlorophyll a/b protein complex. Photosynth. Res. 27, 57–64 (1991).

36. W. I. Gruszecki, P. Kernen, Z. Krupa, R. J. Strasser, Involvement of xanthophyll pigments in regulation of light-driven excitation quenching in light-harvesting complex of Photosystem II. Biochim. Biophys. Acta Bioenerg. 1188, 235–242 (1994).

37. V. Barzda, M. Vengris, L. Valkunas, R. van Grondelle, H. van Amerongen, Generation of fluorescence quenchers from the triplet states of chlorophylls in the major light-harvesting complex II from green plants. Biochemistry 39, 10468–10477 (2000).

38. W. Grudziński et al., Conformational rearrangements in light-harvesting complex II accompanying light-induced chlorophyll a fluorescence quenching. Biochim. Biophys. Acta Bioenerg. 1554, 108–117 (2002).

39. G. S. Schlau-Cohen et al., Single-molecule identification of quenched and unquenched states of LHCII. J. Phys. Chem. Lett. 6, 860–867 (2015).

40. G. S. Schlau-Cohen, Q. Wang, J. Southall, R. J. Cogdell, W. E. Moerner, Single-molecule spectroscopy reveals photosynthetic LH2 complexes switch between emissive states. Proc. Natl. Acad. Sci. U.S.A. 110, 10899–10903 (2013).

41. J. Zielonka, B. Kalyanaraman, Hydroethidine- and MitoSOX-derived red fluorescence is not a reliable indicator of intracellular superoxide formation: another inconvenient truth. Free Radic. Biol. Med. 48, 983–1001 (2010).

42. J. M. Gruber et al., Dynamic quenching in single photosystem II supercomplexes. Phys. Chem. Chem. Phys. 18, 25852–25860 (2016).

43. N. Smirnoff, Ascorbate biosynthesis and function in photoprotection. Philos. Trans. R. Soc. B 355, 1455–1464 (2000).

44. T. P. J. Krüger, C. Ilioaia, M. P. Johnson, A. V. Ruban, R. van Grondelle, Disentangling the low-energy states of the major light-harvesting complex of plants and their role in photoprotection. Biochim. Biophys. Acta Bioenerg. 1837, 1027–1038 (2014).

45. P. Manna, T. Davies, M. Hoffmann, M. P. Johnson, G. S. Schlau-Cohen, Membrane-dependent heterogeneity of LHCII characterized using single-molecule spectroscopy. Biophys. J. 120, 3091–3102 (2021).

46. M. Tutkus, J. Chmeliov, D. Rutkauskas, A. V. Ruban, L. Valkunas, Influence of the carotenoid composition on the conformational dynamics of photosynthetic light-harvesting complexes. J. Phys. Chem. Lett. 8, 5898–5906 (2017).

47. T. Kondo et al., Single-molecule spectroscopy of LHCSR1 protein dynamics identifies two distinct states responsible for multi-timescale photosynthetic photoprotection. Nat. Chem. 9, 772–778 (2017).

48. T. Kondo et al., Microsecond and millisecond dynamics in the photosynthetic protein LHCSR1 observed by single-molecule correlation spectroscopy. Proc. Natl. Acad. Sci. U.S.A. 116, 11247–11252 (2019).

49. A. C. Norris, C. Oberg, L. C. Spangler, G. D. Scholes, G. S. Schlau-Cohen, Discovery of multiple light-harvesting states of the photosynthetic protein PE545. J. Am. Chem. Soc. 146, 27373–27381 (2024).

50. D. H. Stewart et al., Identification of histidine 118 in the D1 polypeptide of photosystem II as the axial ligand to chlorophyll Z. Biochemistry 37, 10040–10046 (1998).

51. C. A. Tracewell, A. Cua, D. H. Stewart, D. F. Bocian, G. W. Brudvig, Characterization of carotenoid and chlorophyll photooxidation in photosystem II. Biochemistry 40, 193–203 (2000).

52. C. A. Tracewell, G. W. Brudvig, Multiple redox-active chlorophylls in the secondary electron-transfer pathways of oxygen-evolving photosystem II. Biochemistry 47, 11559–11572 (2008).

53. R. Gańczarczyk, R. Rybakiewicz-Sekita, E. D. Głowacki, Photoinduced ROS production by poly(3-hexylthiophene) (P3HT) is due to autoxidation rather than catalysis. Chemistry – A European Journal e71488 (2026).

