## Supplementary Material for "Oxygen drives a rapid, reversible photoprotective switch in photosynthetic antennae"

**The PDF file includes:**

Materials and Methods

Supplementary Text

Figs. S1 to S7

Tables S1 to S4

References (*54–58*)

### Materials and Methods

Sample preparation

Thylakoid membranes were isolated from fresh *Spinacia oleracea* following the protocol in ref. (*54*), with modifications as described in ref. (*55*). LHCII trimers were isolated and purified from thylakoid membranes by detergent solubilization followed by sucrose-gradient ultracentrifugation, containing 10 mM HEPES (pH 7.5) and 0.03% (w/v) n-dodecyl α-D-maltopyranoside (α-DM), as described before (*55*). LH2 complexes were isolated from *Rhodoblastus acidophilus* in 12 mM Tris-HCl (pH 8.0) and 0.12% (w/v) lauryldimethylamine oxide (LDAO), as described before (*56*). The samples were stored at −80 °C.

Intensity-Programmed Kinetic Isolation (IPKI)

A schematic of the IPKI setup is shown in Figure 1A. The output of a supercontinuum laser (SuperK EVO, NKT Photonics, 20 MHz) was spectrally filtered using a SuperK VARIA (NKT Photonics) to transmit wavelengths from 525 nm to 625 nm. The filtered beam was directed through an acousto-optic modulator (AOM; 3080-122, Gooch & Housego) and subsequently expanded to produce a Gaussian beam with a diameter of ≈25 mm. A circular iris with a diameter of ≈5 mm was used to select the central portion of the beam to ensure uniform illumination. This region was imaged onto the sample using a wide-field lens and a 10× air objective, producing an excitation spot with a diameter of ≈150 μm in the sample plane. The spot size was verified by co-imaging the excitation beam with a precision pinhole.

Fluorescence was collected through the same objective and separated from the excitation light using a dichroic beamsplitter (FF649-Di01-25x36, Semrock) and a fluorescence filter (ET665lp, Chroma Technology). The filtered fluorescence was focused onto a single-photon avalanche diode (SPAD; SPCM-AQRH-16, Excelitas). A data-acquisition card (PCIe-7846R, National Instruments) controlled both the AOM and SPAD gating.

The AOM was used to switch the excitation beam between high-intensity preparation periods, off periods, and low-intensity probe pulses. The probe pulses had a 10% duty cycle, a repetition frequency of 10–20 Hz, and a corresponding duty-cycle-averaged photon flux density (PFD) of ≈1,000 μmol $m^{-2}$ $s^{-1}$ for the LHCII and Chl *a* measurements and ≈100 μmol $m^{-2}$ $s^{-1}$ for the thylakoid measurements. Residual AOM leakage during the nominally off periods was ≈50 μmol $m^{-2}$ $s^{-1}$. The SPAD was gated on only during the probe pulses. A total of 3–10 runs were averaged per measurement.

Samples were prepared by placing ≈4 μL of sample on a poly-L-lysine (PLL)-coated coverslip and sandwiching the sample with a second coverslip. LHCII samples were diluted in 20 mM HEPES (pH 7.5) and 0.03% (w/v) α-DM to a concentration of ≈0.05 mg $mL^{-1}$ Chl, whereas thylakoid samples were diluted in 20 mM HEPES (pH 7.5), 0.4 M sorbitol, and 5 mM $MgCl_2$ to a concentration of ≈0.5 mg $mL^{-1}$ Chl. Chl *a* was dissolved at ≈1 μM in a 40% glycerol/60% ethanol solution. For oxygen-scavenging experiments, a mixture of 7.5 mg $mL^{-1}$ glucose, 0.2 mg $mL^{-1}$ glucose oxidase, and 0.035 mg $mL^{-1}$ catalase was used. For experiments with AA, the final concentration was 20 mM. At the experimental pH of 7.5, AA is predominantly present as the ascorbate monoanion. For experiments with SOD, the final concentration was 0.2 mg $mL^{-1}$. All measurements were performed at room temperature.

The intensity-dependent IPKI traces were fitted globally using the kinetic models shown in Figure S1A (inset) for LHCII and Figure 4, A and C (insets) for the thylakoids. These are the simplest models that reproduced all experimental observations without systematic residuals and should be regarded as an effective description of the observed dynamics rather than a unique microscopic representation. Additional complexity was introduced only when necessary to eliminate systematic deviations, thereby enabling robust extraction of the dominant kinetic processes while avoiding over-parameterization. Therefore, besides the introduction of a second fluorescent population ($U_2$) for the LHCII and thylakoid data, a biexponential function significantly improved the fit of the recovery process ($k_2$) for the LHCII data (Fig. S1, A and B), whereas for the thylakoid data, a single exponential provided an adequate fit. For LHCII, the quenched population Q was therefore split into two effective populations, $Q_1$ and $Q_2$, populated in fractions $f$ and $1 - f$ and relaxing with rates $k_{2a}$ and $k_{2b}$, respectively, where $k_{2a}$ denotes the fast component and $k_{2b}$ the slow component. The use of $f$ avoids introducing an additional decay component. In the main text, $k_2$ denotes the corresponding effective recovery rate of the pooled Q population. For simplicity, only the average $k_1$ and $k_2$ values are shown in the main text.

The kinetic equations for the full model are given by

$$\begin{pmatrix} B' \\ Q_1' \\ Q_2' \\ U_2' \\ U_1' \end{pmatrix} = \begin{pmatrix} 0 & 0 & 0 & k_4 & k_3 \\ 0 & -k_{2a} & 0 & 0 & fk_1 \\ 0 & 0 & -k_{2b} & 0 & (1-f)k_1 \\ 0 & 0 & 0 & -k_4 & 0 \\ 0 & k_{2a} & k_{2b} & 0 & -k_1 - k_3 \end{pmatrix} \begin{pmatrix} B \\ Q_1 \\ Q_2 \\ U_2 \\ U_1 \end{pmatrix}$$

The solution to these equations is given by a matrix exponential and was fitted to the experimental traces, separating the high-intensity ('light') and low-intensity ('dark') phases whilst ensuring continuity. For the 'dark' phases, $k_1$, $k_3$, and $k_4$ were set to zero. The recovery rate $k_2$ was determined globally and shared across all excitation conditions. Tests of the light dependence of recovery and the mixed-effects treatment are described in Text S2. Nonlinear least-squares regression was used for fitting.

Immobilization of the complexes eliminated translational diffusion and thereby simplified the kinetic model. Reversible quenching nevertheless persisted for freely diffusing LHCII complexes (Fig. S5D), consistent with previous observations of blinking in freely diffusing LHCII complexes (*39*). These observations support the conclusion that immobilization on PLL did not induce the reversible quenching behavior observed here.

DHE/SOD spectroscopy

Fluorescence spectra were measured using a home-built confocal microscope setup (*57*) equipped with a SuperK EVO laser (NKT Photonics, 20 MHz) as an excitation source, which was spectrally filtered (SuperK VARIA, NKT Photonics). LHCII was used at a concentration of 0.05 mg $mL^{-1}$ Chl, with 0.1 mM dihydroethidium (DHE) and 0.2 mg $mL^{-1}$ SOD. Samples were excited at 405 nm with a peak excitation intensity in the focused laser spot of ≈100 W $cm^{-2}$. Fluorescence was filtered using a dichroic beamsplitter (DMLP550R) and a long-pass filter (FELH0550, Thorlabs), diffracted by a grating (GR25–0608, 600 lines/mm, 750 nm blaze, Thorlabs), and collected by an electron-multiplying charge-coupled device (EMCCD) camera (Andor iXon3). To generate superoxide, the entire sample was illuminated for 4 min at ≈5 W $cm^{-2}$ using a Schott KL1500 lamp after it was sent through a 650-nm long-pass filter (FELH0650, Thorlabs) to minimize direct photobleaching of DHE.

Single-molecule spectroscopy (SMS)

A home-built confocal microscope setup (*57*) was used for SMS measurements. Briefly, the excitation light was provided by a supercontinuum laser (SC400-4-PP, 40 MHz, Fianium). For LHCII measurements, 646 nm was selected using an acousto-optic tunable filter (AOTF, Crystal Technology, Inc.), while a bandpass filter (FBH780-10, Thorlabs) was used to excite LH2 with 780 nm light. The excitation beam was circularly polarized by a quarter-wave plate (Edmund Optics) and directed by a dichroic beamsplitter (T660lpxr, Chroma Technology for LHCII, and FF801-Di02, Semrock for LH2) to the back aperture of an oil-immersion objective (1.4 NA, Nikon 100×), which focused the beam onto the sample positioned on a piezo-driven stage (Mad City Labs, LPS200). The excitation fluence was ≈4.2 μJ $cm^{-2}$ per pulse and ≈30.0 μJ $cm^{-2}$ per pulse for LHCII and LH2, respectively. The fluorescence emission was collected by the same objective and separated from the excitation light using fluorescence filters (ET665lp, Semrock for LHCII, and FELH0850, Thorlabs for LH2). The fluorescence was divided by a 70:30 beam splitter, with 70% sent to a SPAD (SPCM-AQRH-16, Excelitas) and registered by a time-correlated single-photon counting (TCSPC) module (Becker & Hickl SPC-130EM) and 30% dispersed via a diffraction grating (GR25–0608, 600 lines/mm, 750 nm blaze, Thorlabs) onto an EMCCD camera (Andor iXon3) to determine the spectral integrity of the complexes.

Immediately before measurements, an aliquot was thawed on ice and diluted to a 1–10 pM concentration using $N_2$-purged buffers and reagents. LHCII complexes were diluted in a 10 mM HEPES (pH 7.5) buffer containing 0.03% (w/v) α-DM and 1 mM $MgCl_2$, the latter of which promoted the attachment of LHCII complexes onto PLL-treated coverslips. LH2 complexes were diluted in 12 mM Tris-HCl (pH 8.0) containing 0.12% (w/v) LDAO. Approximately 10 μL of the diluted LHCII solution was deposited onto a PLL-coated coverslip inside a custom-built sample cell. For enzymatic oxygen scavenging, 7.5 mg $mL^{-1}$ glucose, 750 μg $mL^{-1}$ glucose oxidase, and 100 μg $mL^{-1}$ catalase were added to the sample buffer. To control the ambient oxygen concentration, the samples were flushed using a homogeneous $O_2/N_2$ gas mixture or pure $N_2$ gas (Afrox, 99.99% purity), as shown in Figure S6. 'Ambient' refers to compressed air with a $O_2/N_2$ mixture representing ambient air conditions. The gas was delivered to the sample through a gas nozzle positioned vertically above the sample at a flow rate of 0.1 L $min^{-1}$. No agitation of the sample solution or bubble formation was observed. The samples were first flushed with the gas for about 2 min to allow sufficient time for protein attachment to PLL, after which ≈30 μL of buffer was added to minimize evaporation during the measurements. Fluorescence measurements were initiated after 3–5 min of continuous gas flushing, and the gas flow was maintained throughout data acquisition. The excitation focus was optimized before every measurement. All measurements were performed at room temperature.

Fluorescence intensity traces were analyzed using *Full SMS* (*58*). Only traces consistent with a single fluorescence complex, as indicated by discrete single-step blinking or bleaching, were included. Partially bleached complexes were excluded by selecting only bright complexes and by truncating intensity traces at the onset of progressive or full degradation. Sections of intensity traces affected by stage movement or focal drift were removed from further analysis. A few of the particularly stable complexes were measured more than once. Complexes were typically measured for 60–70 s per measurement, although a few complexes in the samples containing GOC and/or flushed with pure $N_2$ gas were measured for a few minutes at a time. After data filtering, the number of traces retained for analysis was as follows. LHCII: ambient conditions, 33; 130 μM $O_2$, 20; GOC, 169; 55 μM $O_2$, 119; 16 μM $O_2$, 173; $N_2$, 89; GOC + $N_2$, 21. LH2: GOC, 14; GOC + $N_2$, 141. Intensity change points were identified at a 99% confidence level, after which the resolved levels were grouped into ON and OFF states. This binary analysis captures the dominant reversible dark-state transition used to relate SMS measurements to the IPKI kinetic model. Additional intensity levels were not assigned to specific microscopic states. Their occurrence is consistent with the heterogeneous conformational landscape indicated by the multiple fluorescent populations and multiphasic recovery kinetics observed in the bulk measurements.

EPR spectroscopy

Isolated LHCII (OD≈40 at 675 nm) was transferred to a 4-mm EPR tube under dim light and frozen by immersing the tube in cold isopentane. The sample was subsequently measured by EPR. After the measurement, the sample was thawed and illuminated for ≈1 min at ≈180 mW/cm (*8*) using a 448-nm light-emitting diode. The illuminated sample was immediately snap-frozen in isopentane and measured under the same conditions as the control sample.

EPR spectra were recorded at X-band using a Bruker E580 spectrometer equipped with a Bruker Super High-Q (shqe0511) resonator and an Oxford Instruments ESR900 cryostat. The sample temperature was controlled using a ColdEdge Stinger closed-cycle helium cryocooler. The measurement parameters were as follows: microwave frequency: 9.394 GHz; microwave power: 2.561 mW; modulation amplitude: 16 G; temperature: 15 K; magnetic field sweep width: 3634 G; sweep time: 86 s; conversion time: 40.96 ms; time constant: 81.92 ms. The spectra were the average of four signals. The radical signal spectrum was obtained by subtracting the control spectrum from the spectrum recorded after illumination.

### Supplementary Text

Text S1. Comparison with existing fluorescence methods and the rationale for IPKI

Several fluorescence methods can characterize photophysical dynamics, transient dark states, and photobleaching. Fluorescence correlation spectroscopy (FCS) and its lifetime-resolved variants infer molecular dynamics from fluctuations in steady-state fluorescence (*30, 31*). Shrinking-gate FCS can distinguish forward and backward transitions between emissive and dark states and extract corresponding rate constants (*32*). Transient-state (TRAST) spectroscopy probes the population dynamics of long-lived photophysical states through controlled excitation and readout conditions (*33, 34*), while pulse-induced relaxation measurements prepare a photophysical state and monitor its subsequent relaxation (*35–38*). These approaches provide complementary ways to investigate fluorescent-state dynamics, and none should be regarded as intrinsically incapable of resolving millisecond-to-second processes.

The specific challenge addressed here is to distinguish three contributions that can overlap in an observed fluorescence transient: formation of a reversible dark state during intense excitation, recovery after the excitation is reduced or removed, and irreversible photobleaching that accumulates over repeated illumination cycles. These processes can produce similar decreases in fluorescence during a high-intensity interval, but they have different implications. A fluorescence decrease alone does not establish whether the population has entered a recoverable dark state or has been irreversibly bleached. Likewise, observing fluorescence recovery does not quantify the cumulative irreversible loss occurring alongside it. Resolving these contributions within a common kinetic framework is necessary to assess how reversible dark-state cycling relates to photobleaching.

The limitations of fluctuation-based measurements for this task are conditional rather than absolute. The ability of FCS and related approaches to resolve particular transitions depends on the photon statistics, state populations, brightness contrast, concentration, acquisition conditions, and kinetic model. Under weak excitation, where photon counts may be limited, and in heterogeneous systems with multiple overlapping photophysical processes, it can be difficult to constrain all relevant transitions independently. Similarly, TRAST and pulse-induced relaxation can reveal transient-state dynamics, but the measurement protocol and analysis must be designed explicitly if state preparation, recovery under low excitation, and cumulative bleaching are all to be estimated independently. We therefore do not claim that these other approaches cannot measure slow dark states; rather, IPKI is designed to address the particular combination of kinetic contributions required for the present study.

IPKI extends the pulse-induced relaxation approach by using controlled high-intensity preparation intervals, low-intensity fluorescence readout, and repeated cycles. The intense intervals generate a mixture of reversible dark-state population and irreversible photobleaching. Recovery intervals and weak probe pulses then track the return of fluorescence while limiting further perturbation. The progressive loss of fluorescence over repeated cycles constrains the cumulative bleached population, whereas recovery within a cycle constrains the reversible component. A joint kinetic analysis of these responses allows effective rates for dark-state formation, recovery, and bleaching to be estimated from the same experimental sequence. Analysis across excitation intensities further tests which fitted rates vary with excitation and which do not require an intensity-dependent contribution.

Earlier pulse-induced relaxation studies reported reversible light-induced fluorescence quenching in aggregated LHCII and related lipid-containing LHCII preparations (*35–38*). Oxygen dependence was also reported for the quenching in LHCII aggregates (*37, 38*). Those studies established that light-induced quenching can be reversible and oxygen-sensitive, but they did not resolve the corresponding kinetics in isolated LHCII trimers or establish how the process relates to single-complex blinking and irreversible photobleaching. IPKI makes it possible to investigate these contributions in isolated antenna complexes and to compare their kinetics with those of native thylakoid membranes.

IPKI is a kinetic isolation and analysis strategy, not a chemically specific probe. Its fitted state populations and rate constants are effective parameters that describe the measured fluorescence response. Their assignment to particular molecular species or elementary chemical reactions requires independent evidence or the addition of chemical-specific methods. In this study, we therefore combine IPKI with complementary measurements to constrain the interpretation of the reversible dark state, while the detailed identity and configuration of the proposed charge-separated state remain mechanistic questions.

### Text S2. Tests of the light dependence of recovery

To test whether illumination introduced an additional excitation-dependent contribution to recovery, an extra light-on term was allowed to vary with excitation intensity (Fig. S2). This produced no consistent light dependence of $k_2$, whereas $k_1$, $k_3$, and $k_4$ followed a linear dependence on excitation intensity. For LHCII, a two-sided *t*-test of the slope of the linear regression of $k_2$ on excitation intensity gave $p = 0.09$, providing no statistically significant evidence for a linear light dependence. The light-dependent model also described the data less well than the light-independent model, as indicated by its higher Bayesian information criterion (−104 compared to −118). For thylakoids, the corresponding regressions gave $p = 0.72$ in the absence of AA and $p = 0.29$ in its presence, likewise providing no statistically significant evidence for a linear light dependence. These tests cannot exclude a weak light dependence, but they show that the data do not require one. The final model therefore retained a common, intensity-independent $k_2$ for the recovery phases. A mixed-effects treatment was used for $k_2$, $U_2$, and *f* to account for heterogeneity in the dynamics.

### Supplementary Figures

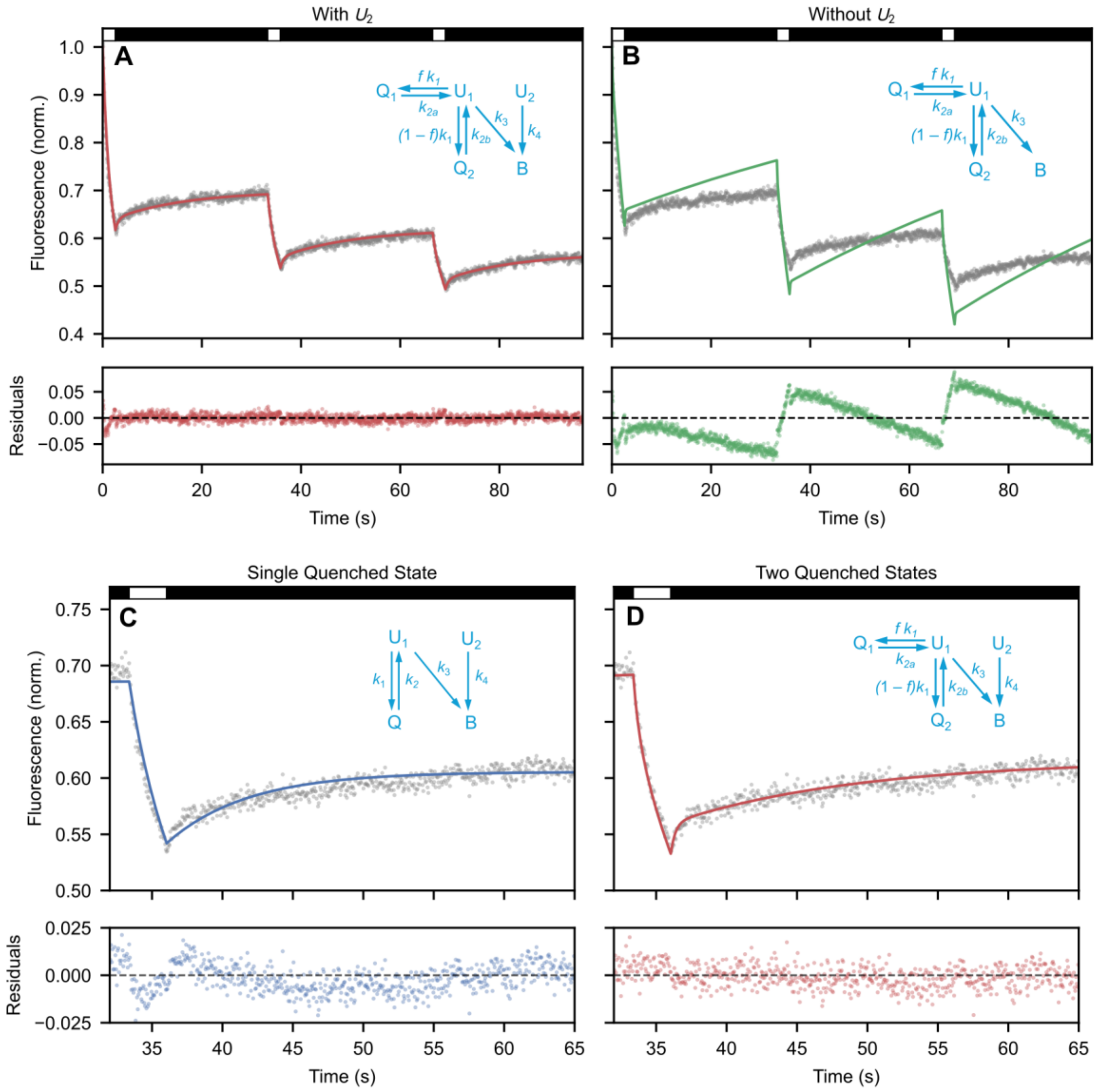


**Fig. S1. Kinetic model refinement**. Top: Need for two effective fluorescent populations. (**A**) Fit (red) of a representative IPKI trace of LHCII (grey) using the full extended kinetic model (inset), which includes both $U_1$ and $U_2$ populations. (**B**) Fit (green) of the same IPKI trace using a kinetic model (inset) that includes only one fluorescent population ($U_1$). Biphasic recovery of Q was used in both cases. Bottom: need for biphasic recovery. (**C**) Fit (blue) of a representative IPKI trace of LHCII (gray) using a kinetic model (inset) that assumes monophasic recovery of Q. (**D**) Fit (red) of the same IPKI trace using a kinetic model (inset) that includes biphasic recovery. Residuals are shown below the fits. Black and white bars on top indicate low- and high-illumination intensities, respectively.

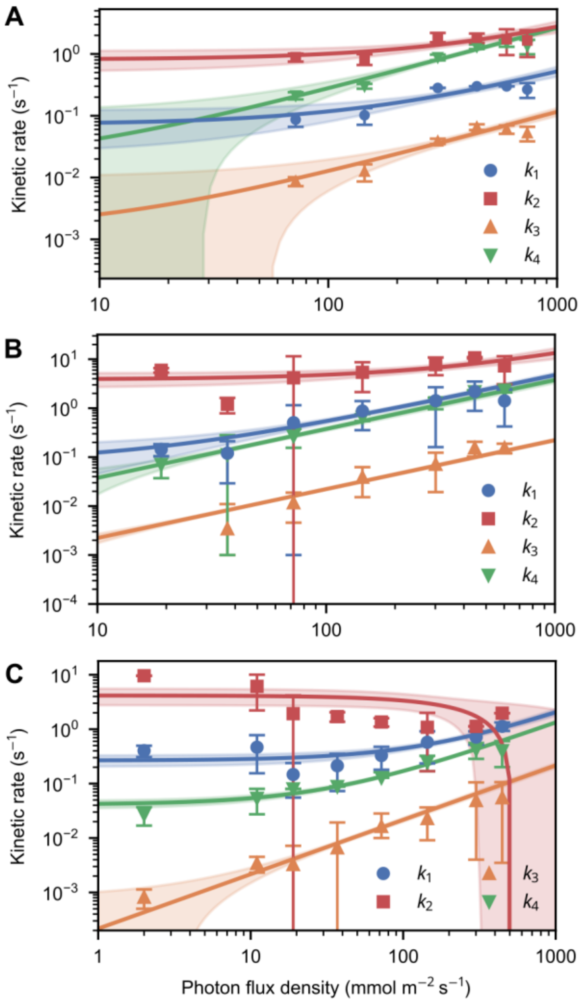


**Fig. S2. Power dependence of IPKI rates with an intensity-dependent light-on recovery contribution.** (**A**), (**B**), and (**C**) correspond to Figures 1A, 4A, and 4C, respectively, but with $k_2$ explicitly allowed to vary with excitation intensity during the high-illumination ("on") periods.

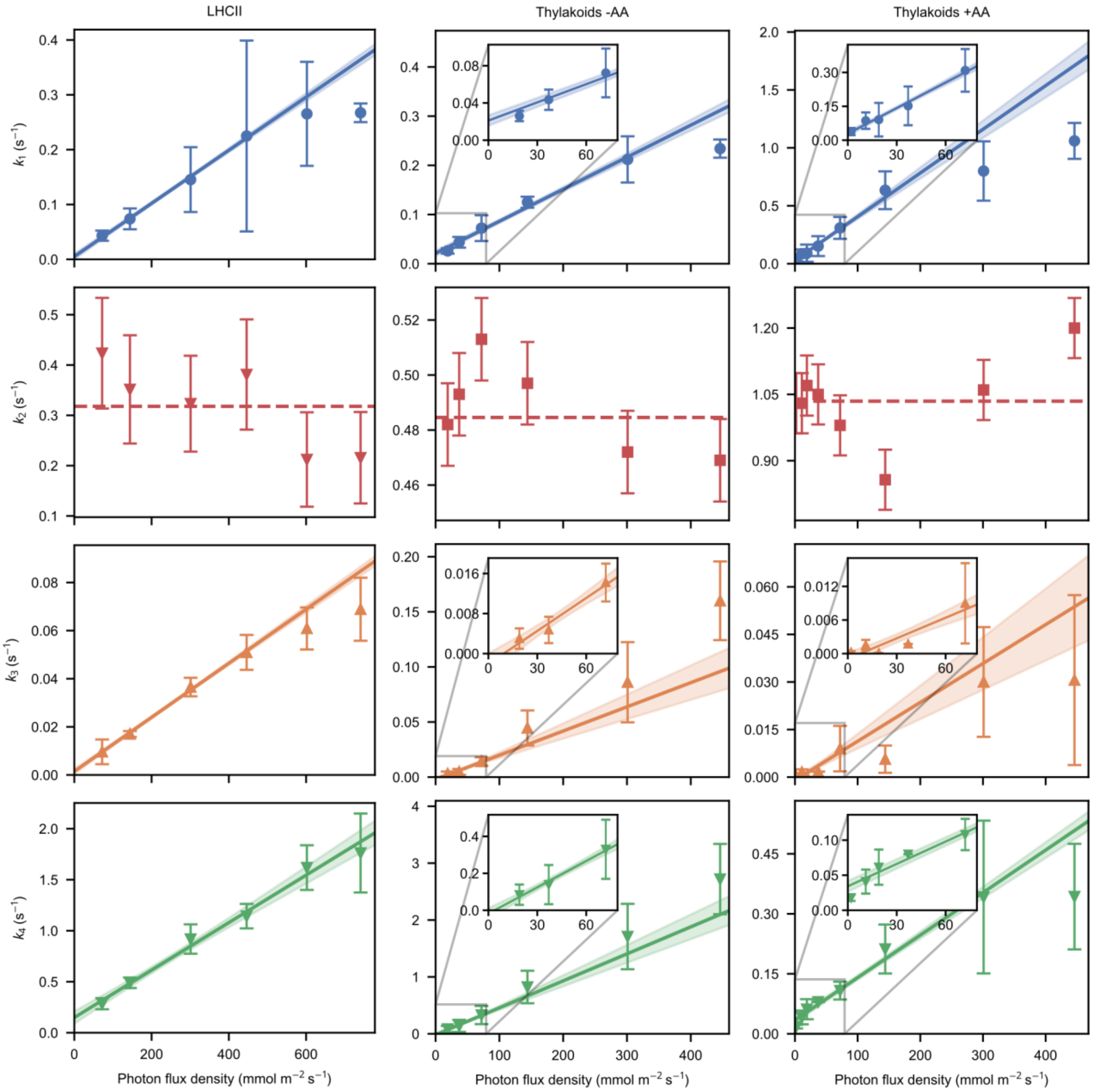


**Fig. S3. Power dependence of kinetic rates on a linear scale. Left**: Same data as for Figure 1C plotted on a linear scale. **Middle**: Same data as for Figure 4A plotted on a linear scale. **Right**: Same data as for Figure 4C plotted on a linear scale. Data points at the highest PFDs were affected by singlet-triplet annihilation. The variation in $k_2$ is due to the mixed-effects model used during the fit, which allows for small variations in constant parameters.

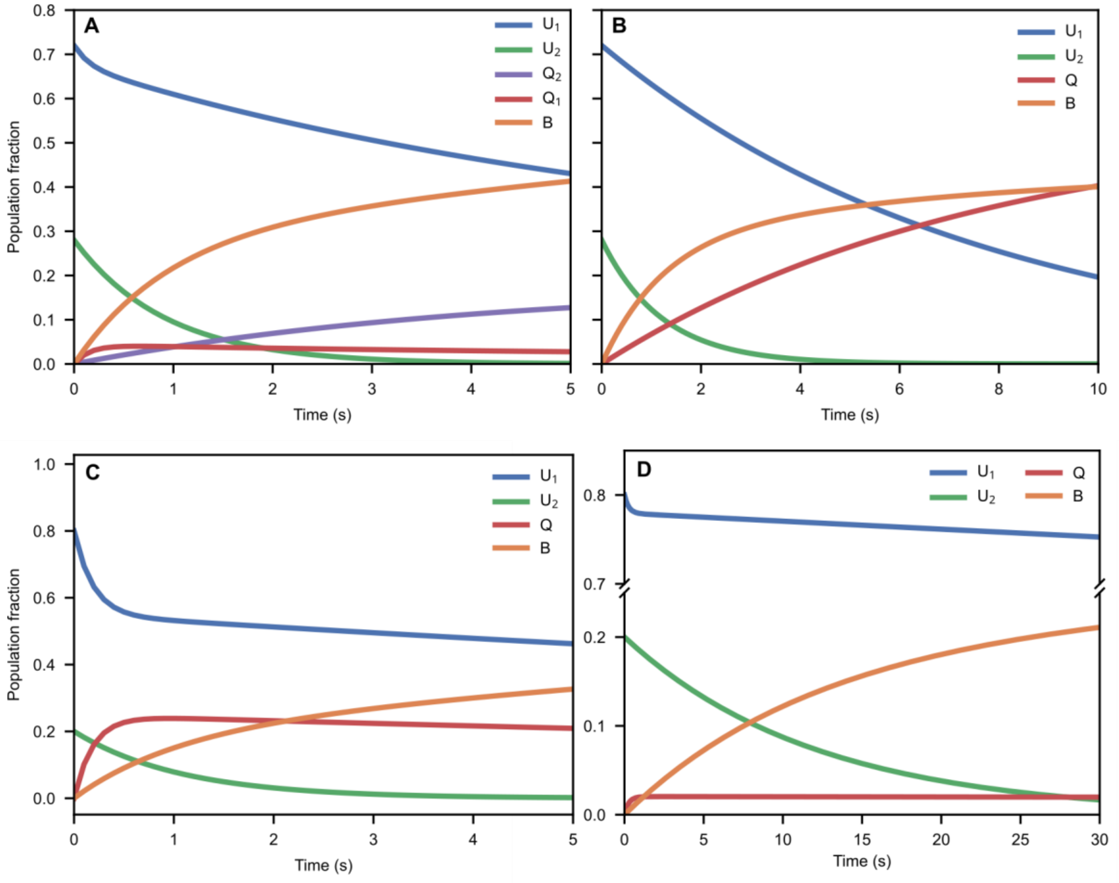


**Fig. S4: Population kinetics for LHCII's extended model and for all samples' accumulated Q**. (**A**) Same as for Figure 1D but showing $Q_1$ and $Q_2$ separately. (**B**) Same as for Figure 1D, but neglecting $k_2$, i.e., considering no fast recovery of Q but its accumulated population over time. The accumulated Q population overtakes the bleached population just before 10 s. (**C**) Simulated population kinetics for thylakoids in the presence of 20 mM ascorbic acid, illuminated continuously with 30,000 μmol photons $m^{-2}$ $s^{-1}$. (**D**) Same as (**C**) but with a PFD of 2,000 μmol $m^{-2}$ $s^{-1}$.

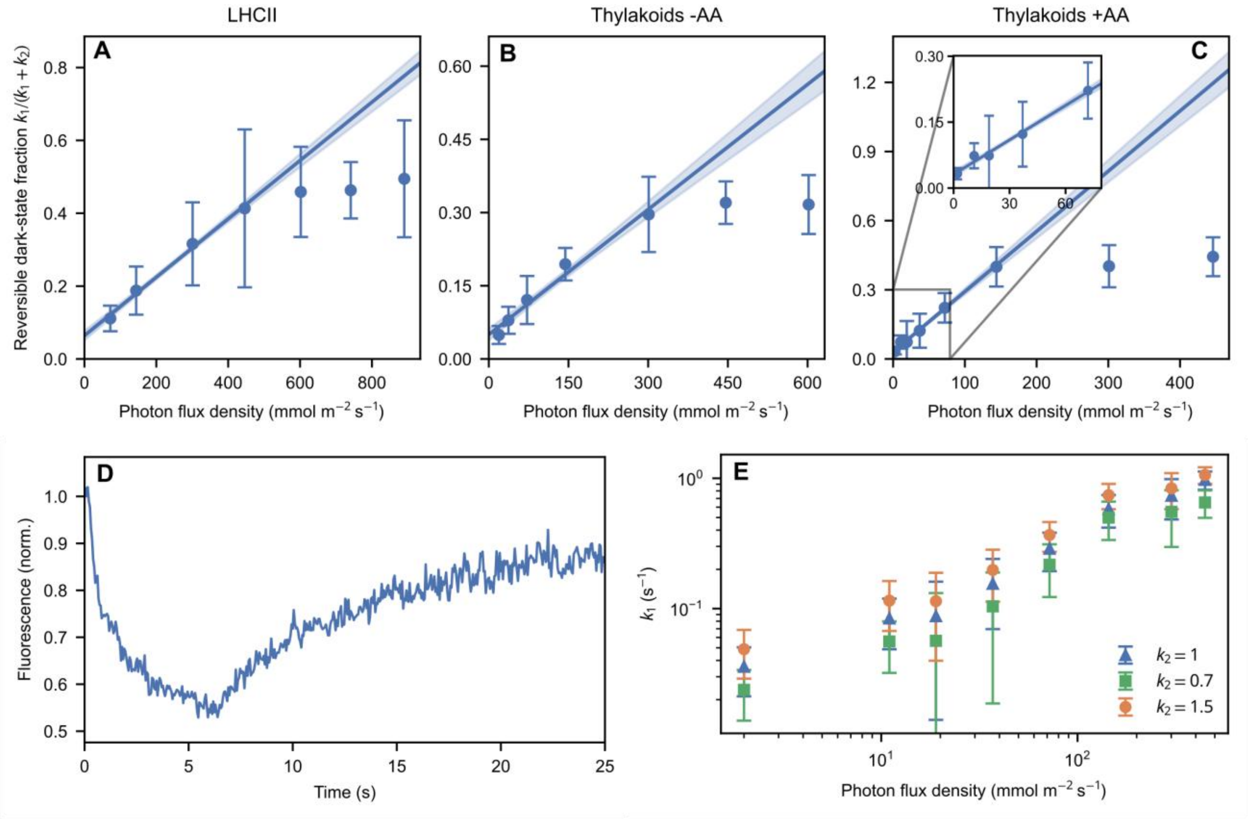


**Fig. S5: Top: Intensity dependence of the reversible dark-state occupancy.** (**A**)–(**C**) Data points were fitted with a linear function (line), with the 68% uncertainty interval shown (shaded). Errors denote Jackknife 95% confidence intervals. For all three samples, the dark-state fraction is linearly correlated with the excitation intensity at low and mid-range intensities, indicating that Q remained far from saturation for these intensities. (**D**) IPKI trace of freely diffusing LHCII, showing that immobilization is not necessary for reversible quenching. (**E**) Effect on the fitted value for $k_1$ when $k_2$ is fixed above (orange) or below (green) its optimal value (blue), for thylakoids with AA. Only a small shift in the fitted $k_1$ values is observed, indicating that the effect of AA on $k_1$ is reliable. The $k_2$ values are given in $s^{-1}$.

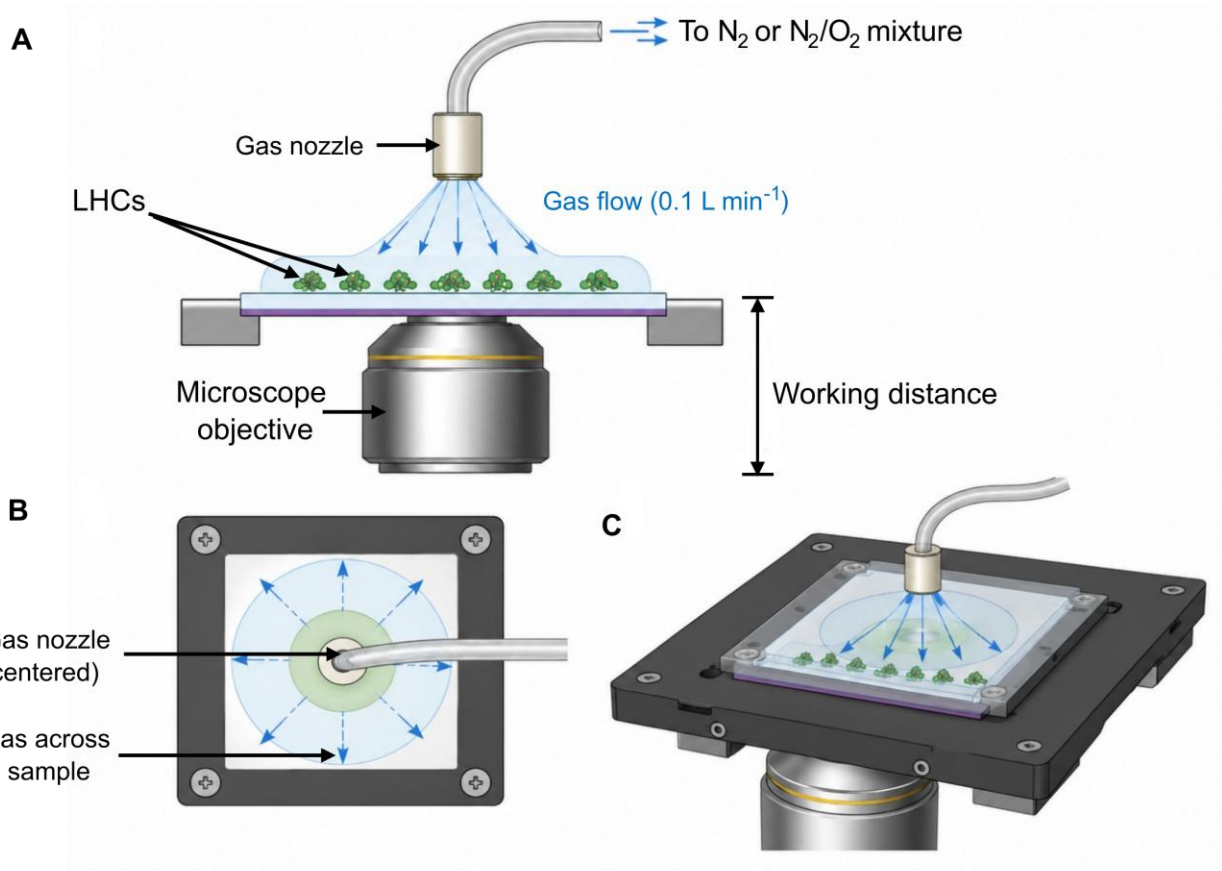


**Fig. S6: Schematic of gas-controlled experimental environment**. (**A**) Side view of the microscope stage showing the gas nozzle positioned above the sample for continuous flushing of $N_2$ or an $O_2$–$N_2$ gas mixture. Not drawn to scale. (**B**) Top view illustrating the gas-flow distribution across the sample surface. (**C**) Three-dimensional representation of the gas-flushing system used during fluorescence measurements.

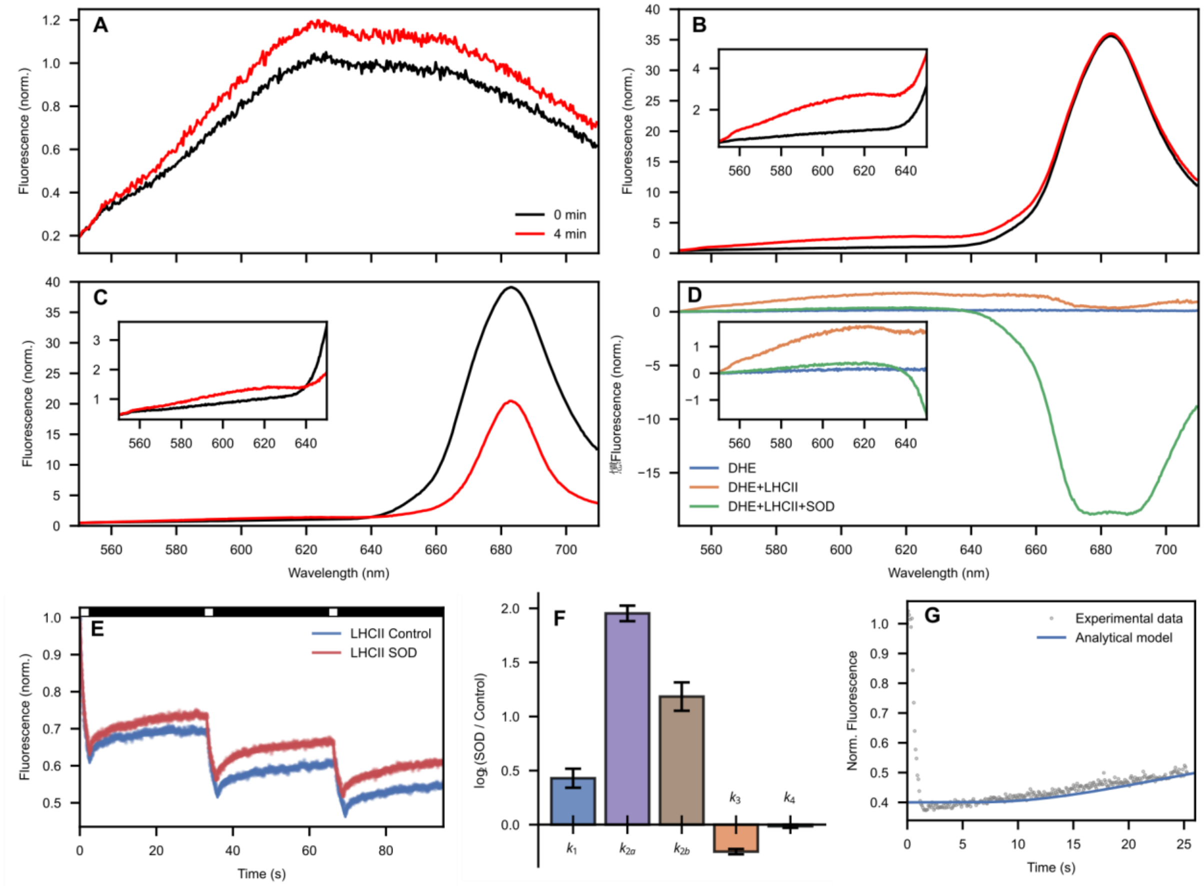


**Fig. S7: Fluorometric study of superoxide formation. (A)** Fluorescence spectrum of a DHE solution under strong external illumination at the onset of illumination (black) and after 4 min of continuous illumination (red). The weak emission upon 405 nm excitation can be attributed to the low probability of DHE to undergo spontaneous and light-induced oxidation, forming mainly 2-hydroxyethidium and ethidium. **(B)** Same as (**A**), but for a solution of LHCII mixed with DHE. Evidently, DHE itself does not measurably accelerate LHCII bleaching under the illumination conditions used. The inset shows a zoom-in around the emission region of the oxidized products of DHE. (**C**) Same as (**B**), but for a solution of LHCII mixed with both DHE and SOD. The significant photobleaching in the presence of SOD is consistent with the removal of superoxide destabilizing the charge-separated state and diverting the Chl cation toward competing irreversible electron-transfer pathways, likely oxidizing nearby molecules. The zoom-in shows considerably less emission of the oxidized products of DHE. **(D)** Same as Figure 3A but extended into the NIR. The DHE spectrum (blue), DHE+LHCII spectrum (orange), and DHE+LHCII+SOD spectrum are the differences between the two spectra in (**A**), (**B**), and (**C**), respectively. The inset shows the same spectra in a narrower wavelength region. (**E**) IPKI trace of LHCII with (red) and without (blue) SOD. The number of measured traces for each sample was 10 (measured on 1 of each sample). **(F)** Same as Figure 3C but showing the values for $k_{2a}$ and $k_{2b}$, which are combined as an effective $k_2$ in Figure 3B. **(G)** Fluorescence recovery after photobleaching (FRAP) of Chl $a$. The experimental data (grey) are the same as for Figure 3D. The theoretical prediction (blue) agrees well with the experimental data, demonstrating that the recovery is due to diffusion and not a long-lived dark state.

### Supplementary Tables

**Table S1:** Effect of oxygen scavengers (GOC) on the IPKI kinetic parameters of LHCII complexes. The branching ratio $f$ was fixed at 0.277, and the initial population of $U_2$ was fixed at 0.25. The number of measured traces for each sample was 10.

| | $k_1$ ($s^{-1}$) | $k_{2a}$ ($s^{-1}$) | $k_{2b}$ ($s^{-1}$) | $k_2$ ($s^{-1}$) | $k_3$ ($s^{-1}$) | $k_4$ ($s^{-1}$) |
|---|---|---|---|---|---|---|
| Ambient conditions | $0.142 \pm 0.004$ | $0.114 \pm 0.004$ | $1.7 \pm 0.1$ | $0.213 \pm 0.009$ | $0.0523 \pm 0.0002$ | $1.39 \pm 0.02$ |
| GOC addition | $0.035 \pm 0.005$ | $0.18 \pm 0.02$ | $1.5 \pm 0.5$ | $0.33 \pm 0.04$ | $0.0013 \pm 0.0001$ | $0.0985 \pm 0.0008$ |

**Table S2:** LHCII's IPKI kinetic parameters corresponding to Figure 1C. PFD: photon flux density. $U_{2,i}$: initial $U_2$ population fraction. $n$: number of individually measured traces for each power level. The branching ratio between $Q_1$ and $Q_2$ was $f = 0.277 \pm 0.001$.

| PFD (mmol $m^{-2}$ $s^{-1}$) | $k_1$ ($s^{-1}$) | $k_{2a}$ ($s^{-1}$) | $k_{2b}$ ($s^{-1}$) | $k_2$ ($s^{-1}$) | $k_3$ ($s^{-1}$) | $k_4$($s^{-1}$) | $U_{2,i}$ | $n$ |
|---|---|---|---|---|---|---|---|---|
| 72 | 0.046 ± 0.009 | 0.14 ± 0.03 | 3.201 ± 0.008 | 0.46 ± 0.08 | 0.010 ± 0.005 | 0.28 ± 0.05 | 0.24 ± 0.04 | 11 |
| 144 | 0.08 ± 0.02 | 0.11 ± 0.03 | 3.199 ± 0.008 | 0.37 ± 0.08 | 0.017 ± 0.001 | 0.49 ± 0.05 | 0.19 ± 0.04 | 11 |
| 301 | 0.15 ± 0.06 | 0.10 ± 0.03 | 3.198 ± 0.008 | 0.34 ± 0.08 | 0.037 ± 0.004 | 0.9 ± 0.1 | 0.28 ± 0.04 | 10 |
| 446 | 0.19 ± 0.18 | 0.12 ± 0.03 | 3.201 ± 0.008 | 0.40 ± 0.08 | 0.051 ± 0.007 | 1.1 ± 0.1 | 0.31 ± 0.04 | 10 |
| 602 | 0.28 ± 0.09 | 0.07 ± 0.03 | 3.214 ± 0.008 | 0.22 ± 0.08 | 0.061 ± 0.009 | 1.6 ± 0.2 | 0.25 ± 0.04 | 10 |
| 741 | 0.28 ± 0.02 | 0.07 ± 0.03 | 3.220 ± 0.008 | 0.23 ± 0.08 | 0.069 ± 0.01 | 1.8 ± 0.4 | 0.20 ± 0.04 | 11 |

**Table S3:** IPKI kinetic parameters of thylakoids in the absence of ascorbic acid, corresponding to Figure 4A. PFD: photon flux density. $U_{2,i}$: initial $U_2$ population fraction. $n$: number of individually measured traces for each power level.

| PFD (mmol $m^{-2}s^{-1}$) | $k_1$ ($s^{-1}$) | $k_2$ ($s^{-1}$) | $k_3$ ($s^{-1}$) | $k_4$ ($s^{-1}$) | $U_{2,i}$ | $n$ |
|---|---|---|---|---|---|---|
| 19 | $0.028 \pm 0.006$ | $0.54 \pm 0.02$ | $0.003 \pm 0.002$ | $0.09 \pm 0.06$ | $0.19 \pm 0.02$ | 11 |
| 37 | $0.05 \pm 0.01$ | $0.55 \pm 0.02$ | $0.005 \pm 0.003$ | $0.1 \pm 0.1$ | $0.22 \pm 0.02$ | 10 |
| 72 | $0.08 \pm 0.03$ | $0.57 \pm 0.02$ | $0.015 \pm 0.004$ | $0.3 \pm 0.2$ | $0.22 \pm 0.02$ | 9 |
| 144 | $0.13 \pm 0.01$ | $0.55 \pm 0.02$ | $0.05 \pm 0.01$ | $0.9 \pm 0.3$ | $0.23 \pm 0.02$ | 5 |
| 301 | $0.22 \pm 0.05$ | $0.53 \pm 0.02$ | $0.09 \pm 0.04$ | $1.8 \pm 0.6$ | $0.23 \pm 0.02$ | 6 |
| 446 | $0.25 \pm 0.02$ | $0.52 \pm 0.02$ | $0.16 \pm 0.04$ | $2.8 \pm 0.6$ | $0.28 \pm 0.02$ | 6 |
| 602 | $0.24 \pm 0.05$ | $0.51 \pm 0.02$ | $0.16 \pm 0.04$ | $3.0 \pm 0.9$ | $0.26 \pm 0.02$ | 5 |

**Table S4:** IPKI kinetic parameters of thylakoids in the presence of ascorbic acid, corresponding to Figure 4C. PFD: photon flux density. $U_{2,i}$: initial $U_2$ population fraction. $n$: number of individually measured traces for each power level.

| PFD (mmol m $^{-2}$s $^{-1}$) | $k_1$ (s$^{-1}$) | $k_2$ (s$^{-1}$) | $k_3$ (s$^{-1}$) | $k_4$ (s$^{-1}$) | $U_{2,i}$ | $n$ |
|---|---|---|---|---|---|---|
| 2 | $0.04 \pm 0.01$ | $1.06 \pm 0.07$ | $0.0007 \pm 0.0003$ | $0.024 \pm 0.008$ | $0.213 \pm 0.002$ | 12 |
| 11 | $0.08 \pm 0.05$ | $1.06 \pm 0.07$ | $0.003 \pm 0.001$ | $0.05 \pm 0.02$ | $0.213 \pm 0.002$ | 12 |
| 19 | $0.09 \pm 0.07$ | $1.10 \pm 0.07$ | $0.0024 \pm 0.0001$ | $0.07 \pm 0.03$ | $0.216 \pm 0.002$ | 8 |
| 37 | $0.16 \pm 0.09$ | $1.11 \pm 0.07$ | $0.0050 \pm 0.0001$ | $0.082 \pm 0.001$ | $0.211 \pm 0.002$ | 8 |
| 72 | $0.29 \pm 0.09$ | $1.01 \pm 0.07$ | $0.014 \pm 0.007$ | $0.13 \pm 0.02$ | $0.213 \pm 0.002$ | 10 |
| 144 | $0.58 \pm 0.2$ | $0.87 \pm 0.07$ | $0.015 \pm 0.004$ | $0.25 \pm 0.06$ | $0.219 \pm 0.002$ | 7 |
| 301 | $0.73 \pm 0.3$ | $1.09 \pm 0.07$ | $0.04 \pm 0.02$ | $0.4 \pm 0.2$ | $0.212 \pm 0.002$ | 13 |
| 446 | $0.97 \pm 0.2$ | $1.22 \pm 0.07$ | $0.05 \pm 0.03$ | $0.4 \pm 0.1$ | $0.212 \pm 0.002$ | 10 |

### Supplementary References


54. S. Caffarri, R. Kouřil, S. Kereïche, E. J. Boekema, R. Croce, Functional architecture of higher plant photosystem II supercomplexes. *EMBO J.* **28**, 3052–3063 (2009).

55. P. Xu, L. Tian, M. Kloz, R. Croce, Molecular insights into zeaxanthin-dependent quenching in higher plants. *Sci. Rep.* **5**, 13679 (2015).

56. R. J. Cogdell, A. M. Hawthornthwaite, Preparation, purification, and crystallization of purple bacteria antenna complexes, in *Photosynthetic Reaction Center* (Elsevier, 1993), pp. 23–42. doi:10.1016/B978-0-12-208661-8.50008-5.

57. F. Kyeyune et al., Strong plasmonic fluorescence enhancement of individual plant light-harvesting complexes. *Nanoscale* **11**, 15139–15146 (2019).

58. J. L. Botha, B. Van Heerden, T. P. J. Krüger, Advanced analysis of single-molecule spectroscopic data. *Biophys. Rep.* **4**, 100173 (2024).
